# Osteopontin and iCD8α cells promote intestinal intraepithelial lymphocyte homeostasis

**DOI:** 10.1101/669432

**Authors:** Ali Nazmi, Michael J. Greer, Kristen L. Hoek, M. Blanca Piazuelo, Joern-Hendrik Weitkamp, Danyvid Olivares-Villagómez

## Abstract

Intestinal intraepithelial lymphocytes (IEL) comprise a diverse population of cells residing in the epithelium at the interface between the intestinal lumen and the sterile environment of the lamina propria. Because of this anatomical location, IEL are considered critical components of intestinal immune responses. Indeed, IEL are involved in many different immunological processes ranging from pathogen control to tissue stability. However, despite their critical importance in mucosal immune responses, very little is known about the homeostasis of different IEL subpopulations. The phosphoprotein osteopontin is important for critical physiological processes, including cellular immune responses such as survival of Th17 cells and homeostasis of NK cells, among others. Because of its impact in the immune system, we investigated the role of osteopontin in the homeostasis of IEL. Here, we report that mice deficient in the expression of osteopontin exhibit reduced numbers of the IEL subpopulations TCRγδ^+^, TCRβ^+^CD4^+^, TCRβ^+^CD4^+^CD8α^+^ and TCRβ^+^CD8αα^+^ cells in comparison to wild-type mice. For some IEL subpopulations the decrease in cells numbers could be attributed to apoptosis and reduced cell division. Moreover, we show *in vitro* that exogenous osteopontin stimulates the survival of murine IEL subpopulations and unfractionated IEL derived from human intestines, an effect mediated by CD44, a known osteopontin receptor. We also show that iCD8α IEL, but not TCRγδ^+^ IEL, TCRβ^+^ IEL or intestinal epithelial cells, can promote survival of different IEL populations via osteopontin, indicating an important role for iCD8α cells in the homeostasis of IEL.

**Key Points:**

1. Osteopontin promotes homeostasis of mouse and human IEL, mediated by its ligand CD44
2. iCD8α cells produce osteopontin which impacts the survival of other IEL
3. Lack of osteopontin renders mice susceptible to intestinal inflammation

## Introduction

One of the largest immunological compartments in the body is comprised of intraepithelial lymphocytes (IEL), a group of immune cells interspaced between the monolayer of intestinal epithelial cells (IEC). IEL can be divided into two groups based on T cell receptor (TCR) expression (1-3). TCR^+^ IEL express αβ or γδ chains. TCRαβ^+^ IEL can be further subdivided into TCRαβ^+^CD4^+^, TCRαβ^+^CD4^+^CD8αα^+^, CRαβ^+^CD8αα^+^, and TCRαβ^+^CD8αβ^+^ cells. TCR^neg^ IEL comprise innate lymphoid cells (ILC) (4-6) and lymphocytes characterized by expression of intracellular CD3γ chains (iCD3^+^), some of which express CD8αα (iCD8α cells) (7, 8).

Because of their anatomical location, IEL function as sentinels between the antigenic contents of the intestinal lumen and the sterile environment under the basal membrane of the epithelium. Indeed, TCRγδ IEL surveil for pathogens (9), secrete antimicrobials conferring protection against pathobionts (10), and protect from intestinal inflammation (11). Other IEL, like conventional CD8 T cells that migrate into the epithelium, can protect against *Toxoplasma* infection (12) and reside in this organ as memory cells (13, 14). TCRαβ^+^CD4^+^CD8αα^+^ IEL can prevent development of disease in the T cell adoptive transfer model of colitis (15). iCD8α cells confer protection against *Citrobacter rodentium* infection and may protect against necrotizing enterocolitis in neonates (8), but these cells can also promote intestinal inflammation in some experimental conditions (16). iCD3^+^ IEL are involved in malignances associated with celiac disease (7).

Osteopontin is a glycosylated phosphoprotein encoded by the Spp-1 (secreted phosphoprotein) gene, originally characterized as part of the rat bone matrix (17, 18). Osteopontin is a versatile molecule involved in many physiological and disease processes (19-21). The role of osteopontin in intestinal inflammation is diverse. For example, Spp-1-deficient mice present with milder disease in the trinitrobenzene sulphonic acid and DSS models of colitis (22, 23). In humans with inflammatory bowel diseases (IBD), plasma osteopontin is significantly increased compared to healthy individuals (24, 25). Some reports indicate that osteopontin is downregulated in the mucosa of Crohn’s disease (CD) patients (26), whereas other groups have reported higher osteopontin expression in the intestines of individuals with CD and ulcerative colitis (UC) compared with healthy controls (25, 27). Because of its involvement in IBD, this molecule could be a potential biomarker (28) and has been explored as a therapeutic target in clinical trials (29). These reports clearly underscore the importance of osteopontin in intestinal inflammation and warrant further investigation of this molecule in mucosal immune responses.

Studies of osteopontin in the immune system have provided important insight into the role of this molecule. For example, osteopontin is involved in macrophage chemotaxis (30), inhibition of NK cell apoptosis and promotion of NK cell responses (31), as well as modulation of dendritic cell function (32). In terms of T cells, osteopontin has been shown to stimulate the survival of concanavalin A-activated lymph node T cells *in vitro*, to promote the survival of anti-myelin specific T cells in the central nervous system (33), and to stimulate Th1 responses (34). However, because IEL are different from “conventional” T cells, and have varied and distinct developmental pathways and functions specialized for the environment of the intestinal epithelium (3), what has been investigated for T cells in other anatomical compartments may not directly translate to IEL, underscoring the need to investigate the role of osteopontin in this peculiar immune site. What we know about the role of osteopontin and IEL homeostasis is very limited. For example, it was reported that the frequency and numbers of TCRγδ^+^ IEL were reduced in osteopontin-deficient mice, while unfractionated TCRαβ^+^ IEL numbers remained similar in comparison to wild type controls (35). However, *in vitro* neutralization of IEL-derived osteopontin resulted in decreased survival of TCRγδ and TCRαβ IEL (35), confounding the *in vivo* results. Our group has recently shown that iCD8α IEL enhance the survival of ILC1-like IEL, via osteopontin, impacting the development of intestinal inflammation (36). Here, we hypothesize that osteopontin and iCD8α cells are key components involved in the homeostasis of most IEL populations. In the present report, we investigated this hypothesis by carefully studying the role of osteopontin in the homeostasis of different IEL subpopulations in mice and total IEL derived from human tissue. We present data showing that osteopontin differentially influences the survival, proliferation and migration of distinct IEL subpopulations, and that these effects are mediated in part by one of the many osteopontin ligands, CD44. Furthermore, we show that IEL survival is mediated primarily by iCD8α cell-derived osteopontin, whereas other TCRγδ^+^ and TCRβ^+^ IEL do not contribute, at least *in vitro*, to the survival of IEL. Moreover, our *in vivo* experiments show that IEC-derived osteopontin do not seem to promote IEL survival. Finally, we present evidence of the impact of osteopontin in the development of intestinal inflammation.

## Material and methods

### Mice

C57BL/6J were originally purchased from The Jackson Laboratory (000664) and have been maintained and acclimated in our colony for several years. CD44^-/-^ (005085), CD45.1 (002014), RFP-Foxp3 (008374) and Spp-1^-/-^ (004936) mice on the C57BL/6 background were originally purchased from The Jackson Laboratory. Rag-2^-/-^ mice on the C57BL/6 background were provided by Dr. Luc Van Kaer. Spp-1^-/-^, CD44^-/-^, and Rag-2^-/-^ mice were crossed with C57BL/6J wild-type mice to generate heterozygous offspring, and subsequently bred among themselves to generate mutant mice. Vill-Cre^+/-^ mice were provided by the laboratory of Dr. Keith Wilson. Spp-1^fl/fl^ mice in the C57BL/6J background were generated by the Vanderbilt Transgenic Mouse/ES Cell Shared Resource following an established protocol (37). Briefly, intron 1/2 and intron 3/4 of the Spp-1 gene were targeted for insertion of LoxP sites by CRISPR/Cas9 ribonucleoproteins (RNP) along with a 1528 bp megamer symmetrical donor ssDNA oligonucleotide (IDT). crRNA sequences were 5’-GTGTGATAACACAGACTCAT-3’ and 5’-AACCAGTACCTTACATGTT-3’. Vill-Cre^+/-^Spp-1^fl/fl^ and Vill-Cre^-/-^Spp-1^fl/fl^ littermate mice were generated by crossing Vill-Cre^+/-^ mice and Spp-1^fl/fl^ mice. Spp-1^-/-^Rag-2^-/-^ mice were generated in our colony by breeding Spp-1^+/-^ with Rag-2^+/-^ mice. Male and female mice were used for all experiments. Mice were maintained in accordance with the Institutional Animal Care and Use Committee at Vanderbilt University.

### Lymphocyte isolation

IEL were isolated by mechanical disruption as previously reported (38). Briefly, after flushing the intestinal contents with cold HBSS and removing excess mucus, the intestines were cut into small pieces (∼1 cm long) and shaken for 45 minutes at 37°C in HBSS supplemented with 5% fetal bovine serum and 2 mM EDTA. Supernatants were recovered and cells isolated using a discontinuous 40/70% Percoll (General Electric) gradient. To obtain lamina propria lymphocytes, intestinal tissue was recovered and digested with collagenase (187.5 U/ml, Sigma) and DNase I (0.6 U /ml, Sigma). Cells were isolated using a discontinuous 40/70% Percoll gradient. Spleen cells were isolated by conventional methods.

### Human samples

The Vanderbilt University Medical Center Institutional Review Board approved sample collection (IRB# 090161 and 190182). Peripheral blood mononuclear cells were isolated by ficoll gradient from unidentified healthy adult volunteers as previously described (39). A pathologist from the Vanderbilt Children’s Hospital provided de-identified fresh intestinal tissue specimens from infants. Isolation of human cells associated with the intestinal epithelium was performed as previously described (40). Briefly, tissue was cut in small ∼1 cm pieces and incubated with slow shaking for 30 minutes at room temperature in HBSS (without calcium and magnesium) supplemented with 5% fetal bovine serum, 5mM EDTA and 1% antibiotic mix (pen-strep-AmphoB; Fisher-Lonza). After incubation, cells in the supernatant were recovered.

### Reagents and flow cytometry

Fluorochrome-coupled anti-mouse CD4 (GK1.5), -CD44 (1M7), - CD45 (30F11), CD45.1 (A20), CD45.2 (104), -CD8α (53–6.7), -CD8β (REA793 or H35-17.2), -TCRβ (H57-597), -TCRγδ (eBioGL3), Ki69 (solA15) and isotype controls were purchased from Thermofisher, BD Biosciences or Tonbo. Annexin V and 7AAD were purchased from BD Biosciences. All staining samples were acquired using BD FACS Canto II or 4-Laser Fortessa Flow cytometers (BD Biosciences) and data was analyzed using FlowJo software (Tree Star). Cell staining was performed following conventional techniques. Manufacturer’s instructions were followed for Annexin V staining; briefly, early apoptotic cells were considered as annexin V^+^7AAD^neg^, late apoptotic cells or undergoing necrosis as annexin V^+^7AAD^+^, and necrotic cells as annexin V^-^7AAD^+^. FACS sorting was performed using a FACSAria III at the Flow Cytometry Shared Resource at VUMC.

### In vitro survival assay

FACS-enriched IEL subpopulations were incubated in a 96-well flat-bottom well plate (Falcon, Fisher Scientific) at a density of 5×10^5^ cells/ml in RPMI containing 10% fetal bovine serum. In some groups culture media was supplemented with recombinant osteopontin (2 μg/ml) (R&D) or anti-CD44 (5 μg/ml) (Thermofisher; clone IM7). Cells were cultured in 5% CO_2_ at 37^°^C. At time 0 and every 24 h, an aliquot from the culture was taken to count live cells using trypan blue to exclude dead cells. Percentage of live cells was calculated in reference to time 0. For human samples, total PBMC or IEL were cultured in the presence or absence of recombinant human osteopontin (2 μg/ml) (R&D) and anti-human-CD44 (5 μg/ml) (Thermofisher; clone IM7).

### Co-culture survival assays

CD45^+^ IEL from Spp-1^-/-^ mice were positively selected using magnetic beads (Miltenyi) and cultured (1×10^5^ cells/well) in a 96-well flat-bottomed well plate in RPMI complemented with 10% fetal bovine serum, penicillin/streptomycin, HEPES, L-glutamine and β-mercaptoethanol. These are “read out” cells. Then, these cells were cultured in the presence (1×10^5^ cells/well) or absence of magnetic bead-enriched (Miltenyi) iCD8α cells from Rag-2^-/-^ mice, or TCRβ^+^ or TCRγδ^+^ IEL from CD45.1 WT mice. For normalization purposes, the purity of these cells was determined by flow cytometry. Anti-osteopontin antibodies (2 μg/ml) (R & D; clone AF808) were added to some wells. After 4 h of incubation, cells were recovered and stained for surface markers, 7AAD and Annexin V. For all experiments, IEL were gated according to their size in a forward versus side scatter. Then, cells were selected by their expression of CD45.2 and gated for the individual subpopulations, followed by analysis of 7AAD incorporation and annexin V staining. Increased survival was determined as 100 - [% of annexin V^+^ read-out cells in co-culture x 100 / % of annexin V^+^ cells cultured alone]. Osteopontin concentration from these cultures was determined by an ELISA kit (R&D) following the manufacturer’s instructions.

### Adoptive transfer of total T cells

Total splenocytes from WT mice were depleted of CD19-positive cells using magnetic beads (Miltenyi). Four to 6 million cells were adoptively transferred (i.p.) into Rag-2^-/-^ or Spp-1^-/-^Rag-2^-/-^ mice (similar results were observed regardless of the number of cells transferred). Starting weight was determined prior to injection. Seven or 28 days later, recipient mice were weighed, sacrificed and donor cells from the intestines analyzed by flow cytometry. In some experiments a segment of the colon was excised and prepared for histological examination. In some experiments CD19-depleted splenocytes from CD44^-/-^ mice were adoptively transferred into Rag-2^-/-^ or Spp-1^-/-^Rag-2^-/-^ mice. Mice were weighed weekly for 4 weeks, and cells and colon analyzed as indicated above. Pathological evaluation of the colons was done in a blind fashion by Dr. Piazuelo, following established parameters (41).

### In vitro and in vivo Foxp3 expression

Lamina propria lymphocytes isolated from RFP-Foxp3 mice were cultured in the presence or absence of recombinant osteopontin and anti-CD44 antibodies as described above. At time 0 and 72 h later, cells were analyzed by flow cytometry to detect RFP expression in live TCR^+^CD4^+^ cells. For *in vivo* experiments, CD4^+^RFP^+^ splenocytes were enriched by FACS and 2 x 10^5^ cells were adoptively transferred i.p. into Rag-2^-/-^ or Spp-1^-/-^Rag-2^-/-^ mice. Eight weeks later, IEL were isolated and RFP expression analyzed in CD45^+^TCRβ^+^CD4^+^ donor-derived cells.

### Transcription profile analysis

For gene expression array, RNA was isolated from FACS-enriched IEL subpopulations from 4 individual WT and Spp-1^-/-^ mice. Samples were prepared for RT^2^ Profiler PCR Array (QIAGEN PAMM-012Z) and analyzed following manufacturer’s instructions. For RNAseq analysis, RNA was isolated from FACS-enriched CD45^+^ IEL derived from WT mice cultured for 24 h in the presence or absence of recombinant osteopontin using the QIAGEN RNeasy micro kit. Sequencing was performed on an Illumina NovaSeq 6000 (2 x 150 base pair, paired-end reads). The tool Salmon (42) was used for quantifying the expression of RNA transcripts. The R project software along with the edgeR method (43) was used for differential expression analysis. For gene set enrichment analysis (GSEA), RNAseq data was ranked according to the t-test statistic. The gene sets curated (C2), GO (C5), immunological signature collection (C7) and hallmarks of cancer (H) of the Molecular Signatures Database (MSigDB) were used for enrichment analysis. GSEA enrichment plots were generated using the GSEA software (44) from the Broad Institute with 1000 permutations.

### Real time PCR for osteopontin expression

RNA from colon tissue was isolated using Trizol following the manufacturer’s instructions. First strand cDNA was synthesized using RT2 first strand kit (Qiagen). Real time PCR was performed using RT2 SYBR Green Mastermix (Qiagen). Osteopontin and GADPH primers were from Qiagen.

### Statistical analysis

Statistical significance between 2 groups was determined using Mann-Whitney U-test. For analysis of 3 groups or more, two-way ANOVA followed by Dunn’s multiple comparison tests were used appropriately. All data was analyzed in GraphPad Prism 7 and shown as mean ± standard error mean (SEM). A *P* value <0.05 was considered significant.

## Results

### Osteopontin deficiency has a differential effect on the homeostasis of individual IEL populations

A previous report has shown that mice deficient in osteopontin (Spp-1^-/-^) have reduced TCRγδ^+^ total cell numbers in the small intestine, while unfractionated TCRαβ^+^ IEL numbers remained the same (35). In the same report, *in vitro* experiments showed that the survival of both of these IEL populations was decreased when cultured in the presence of anti-osteopontin antibodies (35). To further investigate, we analyzed in detail the IEL compartment of WT and Spp-1^-/-^ mice. Fig. 1A depicts the gating strategy used. Our analysis showed a decrease in TCRγδ^+^ small intestine IEL, but contrary to the report by Ito et al (35), we observed that Spp-1^-/-^ mice presented significant reduction in the total numbers of total TCRβ^+^ cells and IEL subpopulations TCRβ^+^CD4^+^, TCRβ^+^CD8α^+^ (which includes CD8αβ^+^ and CD8αα^+^), and TCRβ^+^CD4^+^CD8α^+^ in comparison to cells derived from WT mice (Fig. 1B, top rows). IEL deficiencies were also observed in the colons of Spp-1^-/-^ mice (Fig. 1B bottom rows). TCR^neg^ and iCD8α IEL, presented similar total cell numbers in the small intestine of Spp-1^-/-^ and WT mice but reduced numbers in the colons of Spp-1^-/-^ mice (Fig. 1B). Further subdivision of TCRβ^+^CD8α^+^ IEL based on CD8β expression, showed reduction in small intestine TCRβ^+^CD8αα^+^ and TCRβ^+^CD8αβ^+^ IEL numbers in Spp-1^-/-^ mice (Supplemental Fig. 1A and 1B). In the colon, while the number of TCRβCD8αβ^+^ IEL were similar between WT and Spp-1^-/-^ mice, there was a significant reduction in the total cell numbers of TCRβ^+^CD8αα^+^ IEL in Spp-1^-/-^ mice (Supplemental Fig. 1A and 1B). Interestingly, osteopontin deficiency did not affect spleen T lymphocytes (Fig. 1C) or lamina propria CD19^+^, TCRβ^+^CD4^+^, and TCRβ^+^CD8^+^ cells (Fig. 1D), suggesting that the major influence of this molecule is confined to the intestinal IEL compartment.

**Figure 1.**
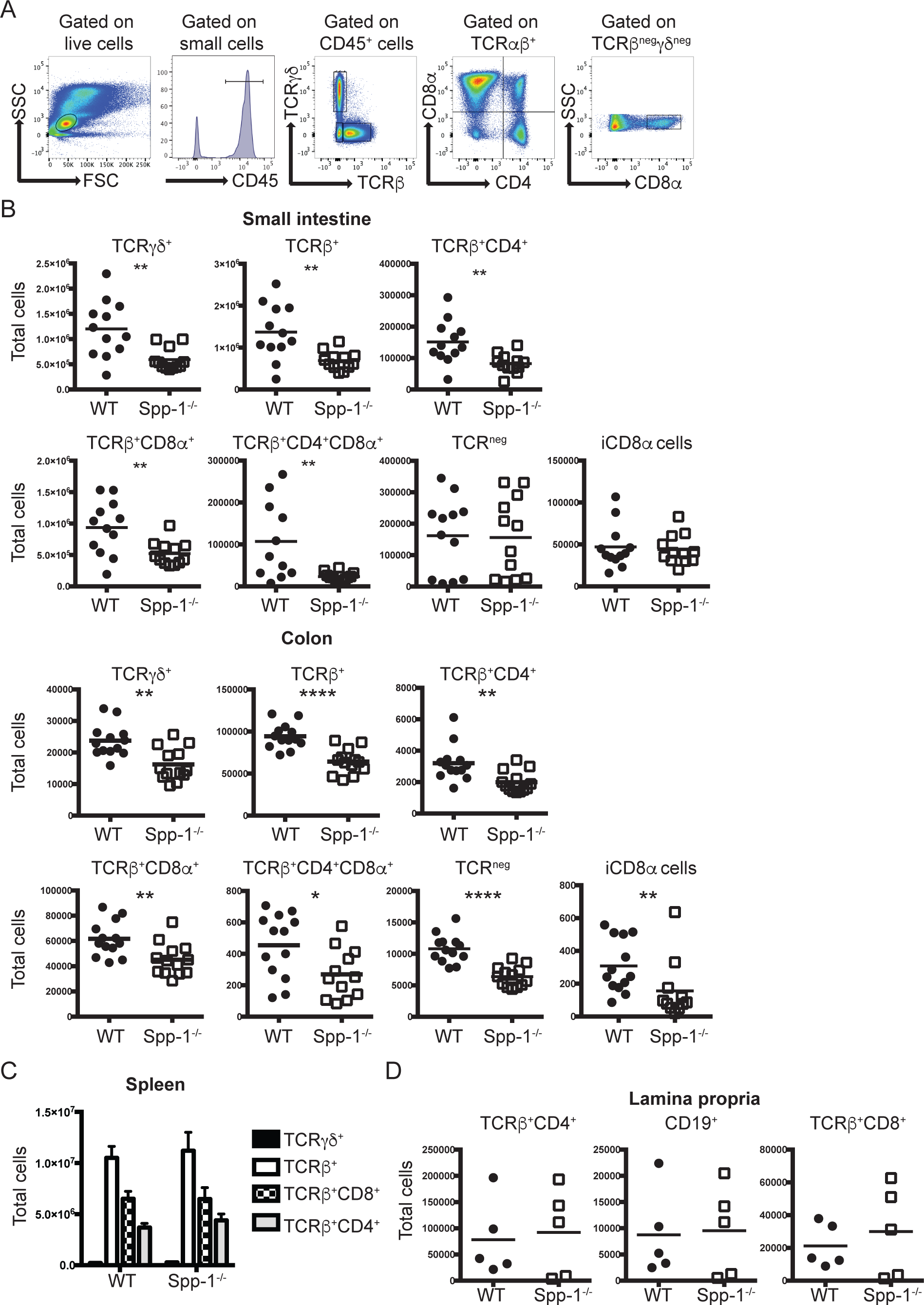
Osteopontin-deficient mice have a reduced IEL compartment. (A) Gating strategy utilized in this report. After gating on live cells, IEL were gated as indicated. (B) Total IEL numbers from the small intestine (top) and colon (bottom) of WT and Spp-1^-/-^ mice. Each dot represents an individual mouse (n = 13). (C) Total cell numbers of the indicated populations in the spleens from WT and Spp-1^-/-^ mice (n = 8). Bars indicate SEM. (D) Total cell numbers of the indicated populations in the lamina propria from WT and Spp-1^-/-^ mice. Each dot represents an individual mouse (n = 5). Data from (B) to (D) are representative of two to three independent experiments. * *P<*0.05; \*\**P*<0.01; \*\*\*\**P*<0.0001 (Mann-Whitney U test).

To investigate whether the reduction in IEL numbers in Spp-1^-/-^ mice was due to increased cell death, we stained the cells with annexin V and 7AAD to determine the levels of early apoptosis (annexin V^+^7AAD^neg^), late apoptosis/undergoing necrosis (annexin V^+^7AAD^+^) and necrosis (annexin V^-^7AAD^+^). TCRγδ^+^, TCRβ^+^CD4^+^, TCRβ^+^CD4^+^CD8α^+^, TCRβ^+^CD8αα^+^, and TCRβ^+^CD8αβ^+^ IEL derived from Spp-1^-/-^ and WT mice presented similar levels of cells in early apoptosis (annexin V^+^7AAD^neg^) and necrosis (annexin V^neg^7AAD^+^) (Fig. 2A). On the other hand, TCRγδ^+^, TCRβ^+^CD4^+^, and TCRβ^+^CD4^+^CD8α^+^ IEL derived from Spp-1^-/-^ mice presented an increase in cells in late apoptosis/undergoing necrosis (annexin V^+^7AAD^+^), suggesting that the decreased total numbers of these IEL subpopulations in the absence of osteopontin may be due to increased cell death.

**Figure 2.**
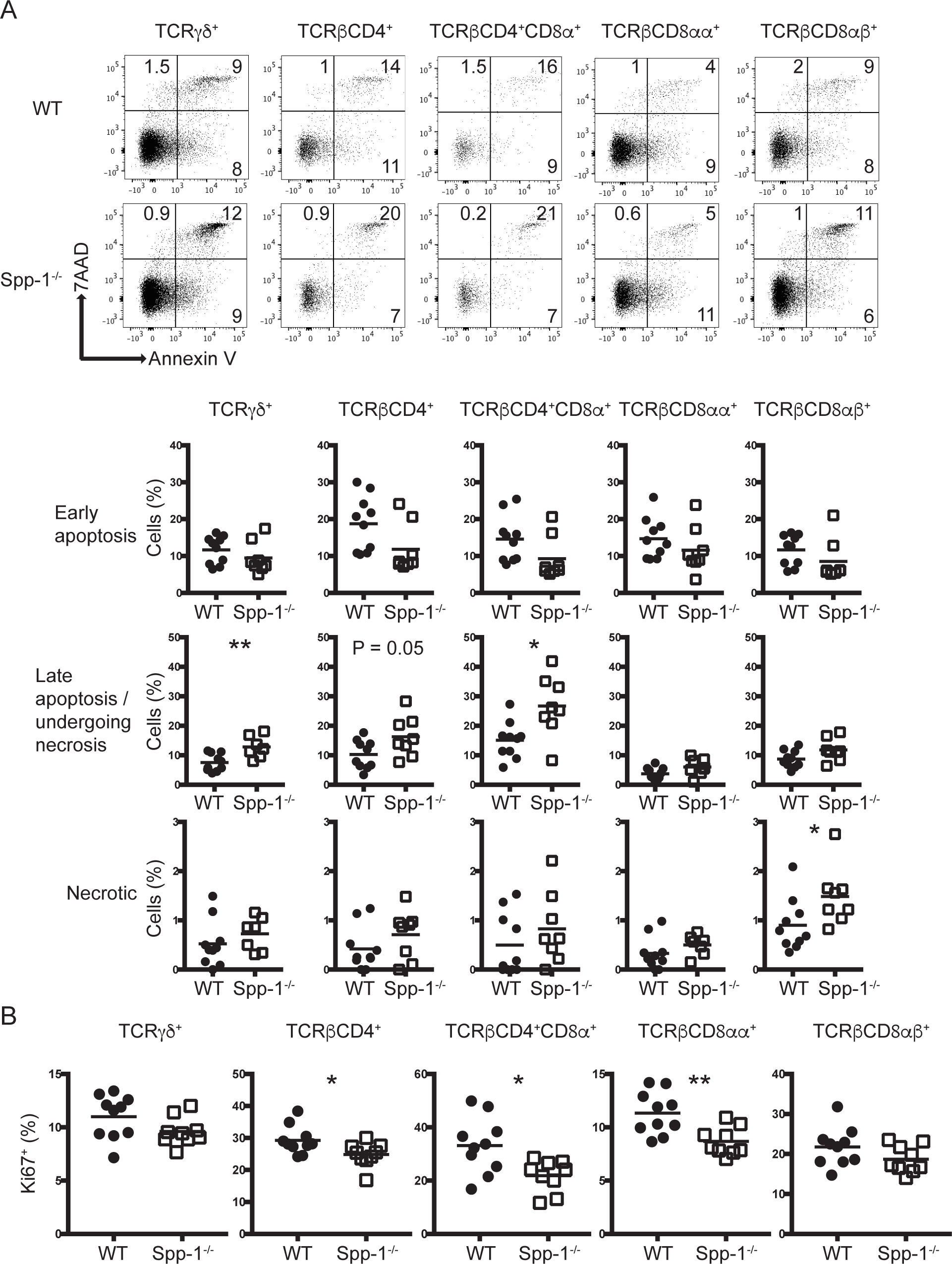
Differential apoptosis and cell division in IEL from osteopontin-deficient mice. (A) Annexin V staining of different small intestine IEL populations derived from WT and Spp-1^-/-^ mice. After gating cells by size based on forward and side scatter profiles, cells were gated for the expression of CD45 and other surface markers as indicated in Fig. 1A, and then analyzed for annexin V and 7AAD staining. Dot plots show a representative sample. Summary is indicated in the graphs. Data are representative of three independent experiments. Each dot represents an individual sample (*n* = 8 to 10). (B) Ki67 intracellular staining of different small intestine IEL populations derived from WT and Spp-1^-/-^ mice. Cells were gated as in Figure 1A. Data are representative from two independent experiments. Each dot represents an individual sample (*n* = 9 to 10). \**P*<0.05; \*\**P*<0.01 (Mann-Whitney U test).

We also analyzed the cell division potential of TCRγδ^+^, TCRβ^+^CD4^+^, TCRβ^+^CD4^+^CD8α^+^ and TCRβ^+^CD8αα^+^ and TCRβ^+^CD8αβ^+^ IEL derived from WT and Spp-1^-/-^ mice. TCRβ^+^CD4^+^, TCRβ^+^CD4^+^CD8α^+^ and TCRβ^+^CD8αα^+^ IEL from osteopontin-deficient mice presented decreased Ki67 staining in comparison to IEL derived from WT mice (Fig. 2B), indicating lower proliferation of these IEL populations. Interestingly, TCRγδ^+^ IEL had similar Ki67 staining levels in cells from osteopontin-deficient and -competent mice (Fig. 2B).

Overall, these results show that osteopontin is required for proper IEL cell numbers, survival and cell division. Importantly, the effect caused by osteopontin is not homogeneous across most IEL populations, and instead, each IEL population responds differently to this cytokine.

### IEL subpopulations survive better in vitro in the presence of osteopontin

To investigate the role of osteopontin in IEL survival, we cultured FACS-enriched TCRγδ^+^, TCRβ^+^CD4^+^, TCRβ^+^CD8α^+^, and TCRβ^+^CD4^+^CD8α^+^ IEL subpopulations in the presence or absence of recombinant osteopontin, and measured survival every 24 h for a period of 3 days. More than 50% of TCRγδ^+^ IEL died after only 24 h of incubation with plain media and continued to die after 48 and 72 h (Fig. 3A). Although TCRγδ^+^ IEL incubated in the presence of osteopontin showed cell death during the first 48 h of cultures, their survival at 72 h was higher than cells incubated without recombinant osteopontin (Fig. 3A). A similar increased survival trend was observed for TCRβ^+^CD8α^+^ and TCRβ^+^CD4^+^CD8α^+^ IEL (Fig. 3A). TCRβ^+^CD4^+^ IEL were more viable than the other populations at 24 h post culture even without recombinant osteopontin, but addition of this cytokine maintained a constant survival rate of TCRβ^+^CD4^+^ IEL (Fig. 3A). Osteopontin binds to several integrin receptors including αvβ1, αvβ3, and αvβ5, among others (45), but also interacts with CD44 (46). Staining TCRγδ^+^ and unfractionated TCRβ^+^ IEL from naïve mice with CD44 results in ∼40 to 50% CD44^+^ cells respectively (Supplemental Fig. 1C). To determine whether the improved survival observed in the presence of osteopontin depends on CD44 binding, IEL subpopulations were incubated in the presence of recombinant osteopontin and anti-CD44 antibody. As shown in Fig. 3A, whenever IEL survival was improved by osteopontin, this effect was blunted by addition of anti-CD44.

**Figure 3.**
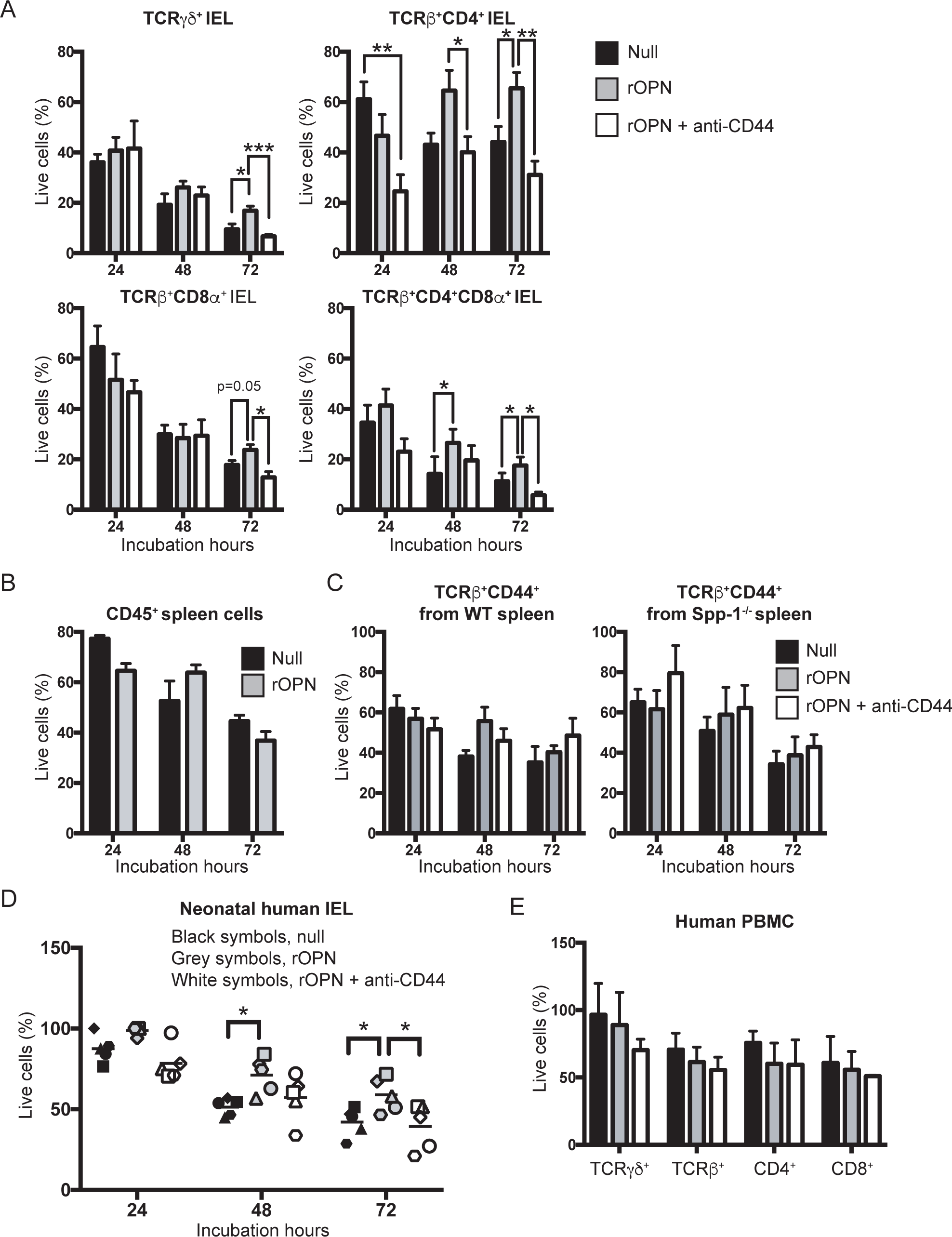
Osteopontin promotes *in vitro* IEL survival. (A) Indicated FACS-enriched IEL populations from WT mice were left untreated or treated with recombinant murine osteopontin (rOPN; 2 μg/ml), with or without anti-mouse CD44 (5 μg/ml). After the indicated time points, cell survival was determined. Data are representative of three independent experiments. Biological replicas consisted of two to three pooled IEL preparations from individual mice; each experiment consisted of 3 biological replicas. (B) Enriched CD45^+^ splenocytes were treated as in (A). Data are representative of two independent experiments (n = 3). (C) FACS-enriched TCRβ^+^CD44^+^ spleen cells from WT and Spp-1^-/-^ mice were treated as in (A). Data are representative of two independent experiments (n = 3). (D) Neonatal human IEL were incubated in the presence or absence of recombinant human osteopontin (2 μg/ml) and anti-human CD44 (5μg/ml). After the indicated time points, cell survival was determined. Each symbol represents an individual human: circle, small intestine from 1-day old patient presenting volvulus and necrosis; square, ileum and colon from 17-day old patient presenting necrotizing enterocolitis; triangle, jejunum from 12-day old patient presenting necrotizing enterocolitis; diamond and hexagon, de-identified individuals. (E) PBMC from adult humans were treated as in (D). Data are representative of two independent experiments (n = 3). \**P*<0.05, \*\**P*<0.01, \*\*\**P*<0.001 (One-way ANOVA).

To determine whether osteopontin-mediated survival is specific for IEL, we investigated the influence of osteopontin on the survival of splenic T cells. First, CD45^+^ splenocytes from naïve WT mice were cultured in the presence or absence of recombinant osteopontin. As shown in Fig. 3B, survival of CD45^+^ spleenocytes was not affected by osteopontin. Moreover, FACS-enriched TCRβ^+^CD44^hi^ spleen cells from naïve WT and Spp-1^-/-^ mice cultured in the presence or absence of recombinant osteopontin and/or anti-CD44 presented no difference in their survival (Fig. 3C), indicating that osteopontin preferentially influences the *in vitro* survival of IEL via CD44 but not of total or CD44^+^ splenic T cells.

The immune system of mice maintained in specific pathogen-free conditions more closely resembles that of human neonates rather than adults (47). Therefore, to determine whether our findings with murine IEL are relevant to humans, we isolated total IEL from human neonates and cultured them in the presence or absence of recombinant human osteopontin. Human IEL survived better in the presence of recombinant osteopontin than in its absence, and the addition of anti-human CD44 blunted the cytokine effect (Fig. 3D), in parallel to the results observed with mouse IEL. To determine the effect of osteopontin on other human lymphocytes, we employed PBMC from healthy adults, as we were unable to obtain PBMCs from the same neonate individuals for this purpose. As shown in Fig. 3E, PBMC survival was not enhanced or reduced by any of the treatments used, which corroborates an intestinal IEL-specific effect. Overall, our results indicate that in *in vitro* conditions, osteopontin differentially promotes both murine and human IEL survival, an effect that is blunted by blocking its interaction with CD44.

Because anti-CD44 blocked the survival effect mediated by osteopontin, we investigated whether the IEL compartment is disrupted in CD44-deficient mice. In comparison to IEL derived from WT mice, CD44^-/-^ mice presented reduced TCR^neg^ and TCRβ^+^CD4^+^ IEL in the small intestines, whereas the colon presented reduction in unfractionated TCRβ^+^, TCR^neg^, TCRβ^+^CD4^+^, and TCRβ^+^CD8α^+^ IEL (Supplementary Fig. 1D).

### Osteopontin induces a differential anti-apoptotic gene expression in IEL

The previous sections suggest an important role for osteopontin in the survival of different IEL subpopulations. To investigate whether osteopontin induces a survival gene expression profile, we isolated RNA from FACS-enriched TCRγδ^+^, TCRβ^+^CD4^+^, TCRβ^+^CD8α^+^ and TCRβ^+^CD4^+^CD8α^+^ IEL derived from naïve WT and Spp-1^-/-^ mice, and determined the expression of genes involved in preventing apoptosis. Comparison of genes expressed in IEL from WT and Spp-1^-/-^ mice showed that TCRγδ^+^ cells from WT animals had more differentially expressed anti-apoptotic genes in comparison to the other IEL populations (Supplemental Fig. 2A and B). TCRβ^+^CD8α^+^ IEL presented little differential expression among the anti-apoptotic genes analyzed. On the other hand, TCRβ^+^CD4^+^ and TCRβ^+^CD4^+^CD8α^+^ IEL differentially expressed some of these genes (Supplemental Fig. 2A and B). Birc2, a known inhibitor of apoptosis in malignancies (48), was one of the genes consistently differentially expressed in most IEL analyzed from WT mice, including TCRβ^+^CD4^+^CD8α^+^ cells (Supplemental Fig. 2A and B). Overall, these results indicate that osteopontin induces the expression of anti-apoptotic genes, but the gene profile varies between different IEL subpopulations.

Because addition of recombinant osteopontin rescued IEL survival when cultured *in vitro* (Fig. 3A), we interrogated whether addition of this cytokine in cultured wild type IEL modifies their gene expression profile. For this purpose, we cultured FACS-enriched CD45^+^ IEL from wild type mice in the presence or absence of osteopontin. Twenty-four hours post-culture, cells were collected, RNA extracted, sequenced and the gene expression profile determined. As recovery of sufficient cells for gene expression profile analysis after 24 h of culture from individual IEL populations was limiting, total CD45^+^ IEL were used as an alternative approach. Gene set enrichment analysis (GSEA) revealed that IEL cultured in the presence of recombinant osteopontin express genes associated with retinoid X receptor (RXR) functions (Supplemental Fig. 2C and D). GSEA also showed that IEL cultured in media alone present enriched pathways related to apoptosis, degradation of p27/p21 and downregulation of genes in regulatory T cells (Supplemental Fig. 2C and D). These results indicate that *in vitro* IEL exposure to osteopontin has an impact on IEL gene transcription and modifies gene sets related to cell survival.

### Osteopontin does not influence T cell migration into the epithelium but maintains homeostasis of TCRβ^+^CD4^+^ and TCRβ^+^CD4^+^CD8α^+^ IEL

Because some IEL populations are derived from conventional CD4^+^ and CD8^+^ T cells, we investigated whether osteopontin is important for the relocation of these cells into the IEL compartment. To test this hypothesis, we adoptively transferred total spleen T cells from WT mice into Rag-2^-/-^ or Spp-1^-/-^Rag-2^-/-^ recipient mice, and after 7 days we determined the number of cells migrating into the intestinal epithelium. Fig. 4A (top) shows the gating strategy to identify cells derived from donor mice. Both TCRβ^+^CD4^+^ and TCRβ^+^CD8α^+^ cells migrated similarly into the epithelium of Rag-2^-/-^ or Spp-1^-/-^Rag-2^-/-^ recipient mice (Fig. 4A, bottom), indicating that osteopontin in the recipient mice does not influence the migration of these cells into the intestinal mucosa. Interestingly, reconstitution analysis of Spp-1^-/-^Rag-2^-/-^ recipient mice at 28 days post transfer showed a reduction in the total number of TCRβ^+^CD4^+^ and TCRβ^+^CD4^+^CD8α^+^ cells but not TCRβ^+^CD8αβ^+^ IEL (Fig. 4B), which resembled what was observed in naïve Spp-1-deficient mice (Fig. 1B and Supplemental Fig. 1B). To prevent the development of intestinal inflammation in Rag-2^-/-^ recipient mice, we transferred total T cells, which includes regulatory T cells. To our surprise, Spp-1^-/-^Rag-2^-/-^ recipient mice lost more weight than Rag-2^-/-^ mice (Fig. 4C) and presented increased colon inflammation (Fig. 4D). The impact of ostopontin produced by donor-derived cells in IEL reconstitution and disease development was minimal, as determined by total colon osteopontin mRNA expression prior to, and after 28 days post T cell transfer (Fig. 4E).

**Figure 4.**
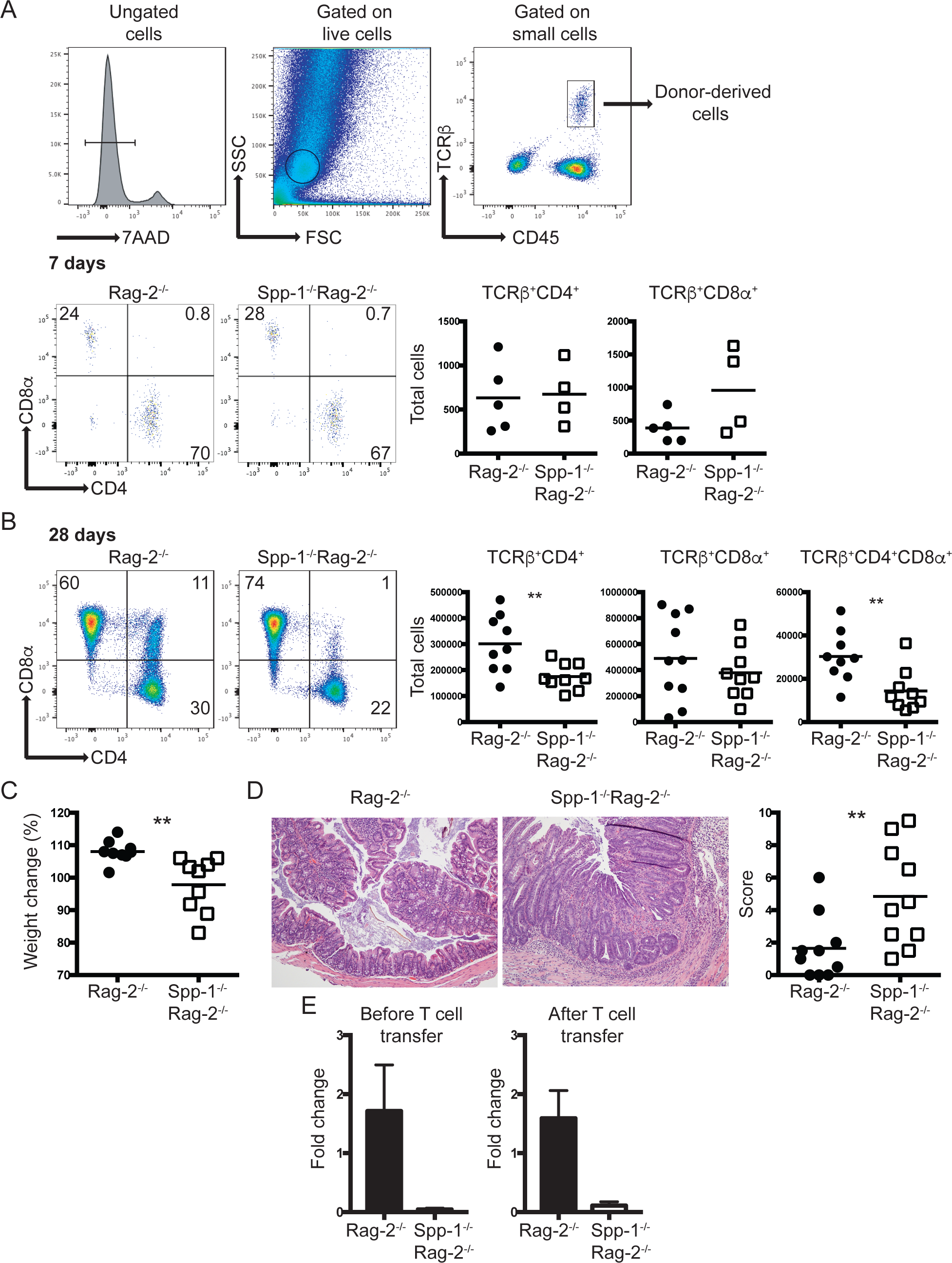
Environmental osteopontin influences IEL reconstitution and colon inflammation. Enriched total spleen T cells from WT mice were adoptively transferred into Rag-2^-/-^ and Spp-1^-/-^ Rag-2^-/-^ mice. (A) Gating strategy to identify cells derived from donor mice (top). Representative analysis of mice analyzed seven days after transfer (bottom). Dot plots show a representative sample. Results are summarized in the graphs. TCRβ^+^CD4^+^CD8α^+^ cells were not detected above background at this time point in either group. Data are representative from two independent experiments. Each dot represents an individual sample (n = 4 to 5). (B) The same analysis as in (A) performed at 28 days post transfer. Dot plots show a representative sample. Results are summarized in the graphs. Data are representative from three independent experiments. Each dot represents an individual animal (n = 9). (C) Total weight change at 28 days post transfer of mice treated as in (B). (D) Representative H&E stained samples of colon sections from the indicated mice at 28 days (100X magnification). Graph indicates total histological score [immune cell infiltration (3 points), loss of goblet cells (3 points), crypt damage (3 points) and epithelial hyperplasia (3 points)]. For (C) and (D) data are representative from three independent experiments, each dot represents an individual mouse (n = 9). (E) Total mRNA expression from colons of naïve and T cell treated (28 days post-transfer) Rag-2^-/-^ and Spp-1^-/-^Rag-2^-/-^ mice (n=3-5). \*\**P*<0.01 (Mann-Whitney T test).

To interrogate why Spp-1^-/-^Rag-2^-/-^ mice developed intestinal inflammation when regulatory T cells were present in the inoculum, we investigated the fate of regulatory T cells in the presence or absence of osteopontin. Regulatory T cells are known to express CD44, which when ligated, promotes sustained Foxp3 expression (49). Thus, we hypothesized that binding of osteopontin to CD44 is a potential signal that maintains proper Foxp3 expression. To test this possibility, we cultured total T cells from the intestines of RFP-Foxp3 mice in the presence or absence of osteopontin, with or without anti-CD44. After 72 h of culture, there was an increase in the percentage of RFP^+^ cells in the presence of osteopontin, which was blunted with the addition of anti-CD44 antibodies (Fig. 5A). Figure 5B shows the combined fold increase over untreated cells.

**Figure 5.**
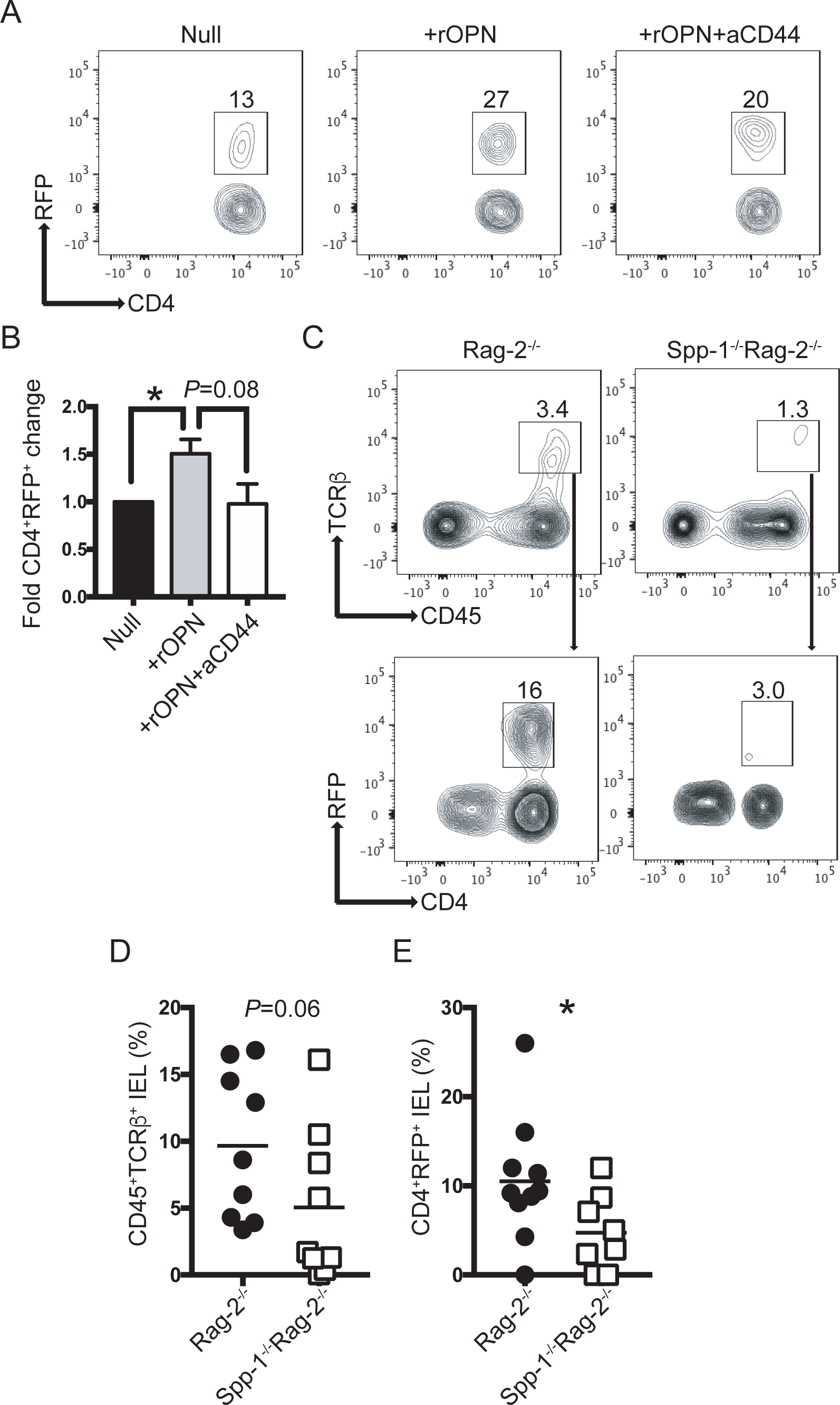
Osteopontin sustains Foxp3 expression. (A) Intestinal T cells from RFP-Foxp3 reporter mice were isolated and cultured in the presence or absence of recombinant osteopontin (2 μg/ml) and anti-CD44 (5 μg/ml). Seventy-two hours later, the percentage of RFP^+^ cells was determined by flow cytometry. After excluding dead cells, dot plots were gated as CD45^+^TCRβ^+^ cells. (B) Bar graph indicates the fold change in CD4^+^RFP^+^ cells in relation to the null group. Data are representative of three independent experiments. (C) Enriched splenic RFP^+^ cells from RFP-Foxp3 reporter mice were adoptively transferred into Rag-2^-/-^ and Spp-1^-/-^Rag-2^-/-^ recipient mice. Eight weeks after transfer, IEL from small intestine were isolated and the percentage of RFP^+^ determined. After excluding dead cells, plots were gated as indicated by the arrows. (D) Summary of the percentage of donor-derived cells recovered; each dot represents an individual mouse. Data are representative from three independent experiments. (E) Summary of the percentage of RFP^+^ cells within the donor-derived cells. \**P*<0.05 (Mann-Whitney T test). rOPN = recombinant osteopontin; aCD44 = anti-CD44 antibodies.

To test whether osteopontin sustains Foxp3 expression *in vivo*, we sorted splenic RFP^+^ cells from RFP-Foxp3 reporter mice and adoptively transferred them into Rag-2^-/-^ or Spp-1^-/-^Rag-2^-/-^ recipient mice. Eight weeks after transfer, IEL were isolated and the percentage of donor-derived (CD45^+^TCRβ^+^CD4^+^) RFP^+^ cells was determined (Fig. 5C). Rag-2^-/-^ mice presented a trend of higher percentage of donor-derived cells in the IEL compartment than Spp-1^-/-^Rag-2^-/-^ recipient mice (Fig. 5D). Approximately 10% of the donor-derived cells from Rag-2^-/-^ recipient mice remained RFP^+^, whereas only 4% of cells recovered from Spp-1^-/-^Rag-2^-/-^ recipient mice remained RFP^+^ (Fig. 5E). These results indicate that osteopontin sustains Foxp3 expression in regulatory T cells in the IEL compartment, possible mediated by CD44 ligation, which significantly impacts the development of intestinal inflammation.

To test whether CD44 expression, as one of the receptors for osteopontin on T cells, is critical for prevention of experimental colitis, we adoptively transferred total T cells from CD44^-/-^ donor mice into Rag-2^-/-^ and Spp-1^-/-^Rag-2^-/-^ recipient mice. Rag-2^-/-^ mice that received total spleen T cells from WT mice did not lose weight, whereas Rag-2^-/-^ and Spp-1^-/-^Rag-2^-/-^ recipient mice that received spleen cells from CD44^-/-^ donor mice lost weight comparably starting at 2 weeks post transfer (Supplemental Fig. 3A), with clear signs of intestinal inflammation (Supplemental Fig. 3B).

Overall, these results show that osteopontin is not involved in the migration of adoptively transferred conventional T cells into the intestinal epithelium, but sustains proper TCRβ^+^CD4^+^ and TCRβ^+^CD4^+^CD8^+^ cell numbers once in the IEL compartment. Moreover, the lack of osteopontin in recipient mice results in development of colitis, even in the presence of regulatory T cells in the inoculum. A similar outcome was observed when CD44-deficient T cells were transferred into Rag-2^-/-^ or Spp-1^-/-^Rag-2^-/-^ recipient mice, highlighting the importance of the osteopontin-CD44 interaction in intestinal homeostasis.

### iCD8α cell-derived osteopontin increases IEL survival

iCD8α cells are a population of TCR^neg^ IEL involved in different immunological roles and are known as an important source of osteopontin in the intestines (8). Recently, it has been shown that survival of TCR^neg^ ILC1-like IEL depends in part on iCD8α cells, an effect most likely associated with the production of osteopontin by these cells (36). To determine whether the survival of TCR^+^ IEL populations is dependent on iCD8α cell-derived osteopontin, we performed experiments in which the source of osteopontin is confined to a specific IEL population (iCD8α, TCRβ^+^ or TCRγδ^+^ cells). Then, these individual IEL populations were co-cultured in the presence of total IEL derived from osteopontin-deficient mice. After 4 h of culture, cells were stained for annexin V and the increased in survival was calculated as indicated in the Materials and method section. As shown in Fig. 6A, addition of osteopontin-competent iCD8α cells to the IEL culture resulted in an increased survival (over IEL cultured in the absence of iCD8α cells) of almost 30% for TCRγδ^+^ and TCRβ^+^CD8αα^+^ IEL, whereas there was variable survival increase for unfractionated TCRβ^+^, and the IEL subpopulations TCRβ^+^CD4^+^, TCRβ^+^CD4^+^CD8α^+^ and TCRβ^+^CD8αβ^+^. Survival was reduced by addition of anti-osteopontin antibodies, indicating that iCD8α cell-derived osteopontin was responsible for the observed increase in survival. A previous report indicated that TCRβ^+^ and TCRγδ^+^ IEL also produce osteopontin (35). To determine whether these IEL populations promote *in vitro* survival of IEL, we performed a similar experiment but the potential source of osteopontin were TCRβ^+^ or TCRγδ^+^ IEL (Fig. 6B and C). TCRβ^+^ IEL were capable of stimulate survival of only TCRγδ^+^ IEL and addition of anti-osteopontin antibodies blunted the survival below the levels of cells cultured alone. For other IEL population, the addition of TCRβ^+^ IEL from osteopontin-competent mice did not enhance IEL survival (Fig. 6B). When TCRγδ^+^ IEL were used as the source of osteopontin, these cells were capable of increasing only the survival of TCRβ^+^CD8αβ^+^ IEL. To determine that the osteopontin-competent cells have similar viability after 4 h of culture, we analyzed their annexin V profile. As indicated in Supplemental Fig. 4A, iCD8α, TCRβ^+^ and TCRγδ^+^ IEL presented low percentage of annexin V-positive cells, and these percentages were similar among the 3 different type of IEL. To determine the production of osteopontin from each cell type in these co-cultures, we recovered the supernatants and measured osteopontin concentration by ELISA. As expected, osteopontin was not detected in supernatants from CD45^+^ IEL derived from Spp-1^-/-^ mice cultured alone, but was readily detectable in cultures containing iCD8α cells (Supplemental Fig. 4A). On the other hand, osteopontin was not detected in the supernatants that included TCRβ^+^ or TCRγδ^+^ IEL from osteopontin-competent mice (Supplemental Fig. 4A). These results suggest that the observed increased survival promoted by TCRβ^+^ and TCRγδ^+^ IEL may not be completely dependent on osteopontin, since this protein was not detected in the supernatants. Overall, these results indicate that iCD8α cell-derived osteopontin is an important survival signal for IEL.

**Figure 6.**
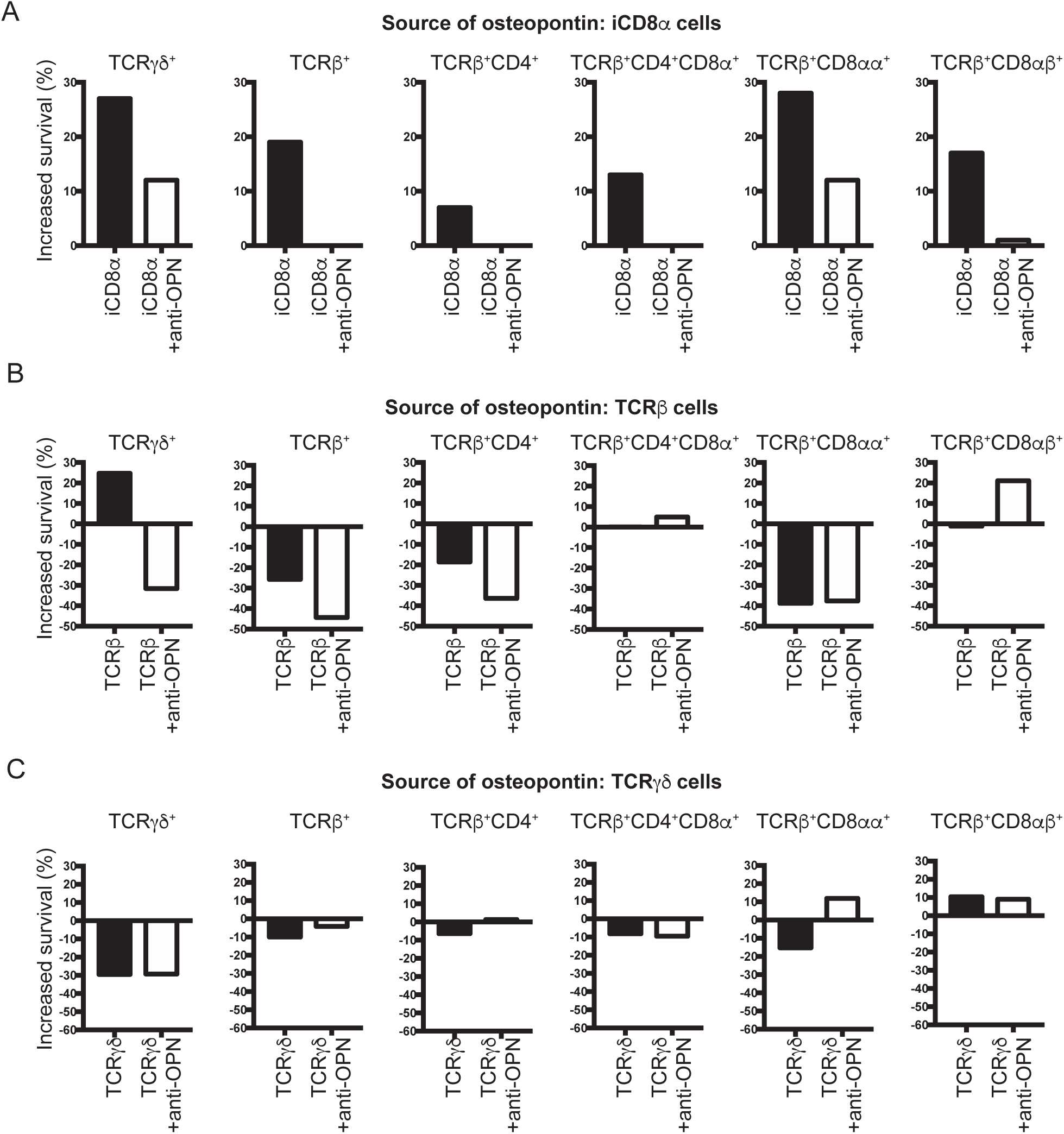
iCD8α cell-derived osteopontin promotes IEL survival. Enriched CD45^+^ IEL from small intestine and colon derived from Spp-1^-/-^ mice were incubated in the presence or absence of enriched iCD8α IEL (A), TCRβ^+^ IEL (B), or TCRγδ^+^ IEL (C) from osteopontin-competent donors. Some cells were cultured in the presence of anti-osteopontin antibodies (2 μg/ml). Four hours after incubation, cells were stained for surface markers, including annexin V, and 7AAD. Cells were gated as in Figure 2A. Graphs indicate “increased survival”, which was determined as 100 - [% of annexin V^+^ read-out cells in co-culture x 100 / % of annexin V^+^ cells cultured alone].

Intestinal epithelial cells are known to constitutively express low levels of osteopontin (25, 26), representing a potential source of this protein for the homeostasis of IEL. Because IEC rapidly die in *in vitro* culture conditions, we determine their contribution to IEL homeostasis by analyzing the IEL compartment of mice with conditional mutation of the osteopontin gene driven by the villin promoter, which is preferentially expressed in IEC (Vill.Cre^+/-^Spp-1^fl/fl^). Expression of osteopontin in IEC is ablated, but not in activated T cells, in Vill.Cre^+/-^Spp-1^fl/fl^ mice validating the conditional mutation (Supplemental Fig. 4C). As shown in Fig. 7A and B, the absence of osteopontin expression in IEC does not impact total small intestine or colon IEL cell numbers, indicating that IEC-derived osteopontin does not contribute to the homeostasis of most IEL populations.

**Figure 7.**
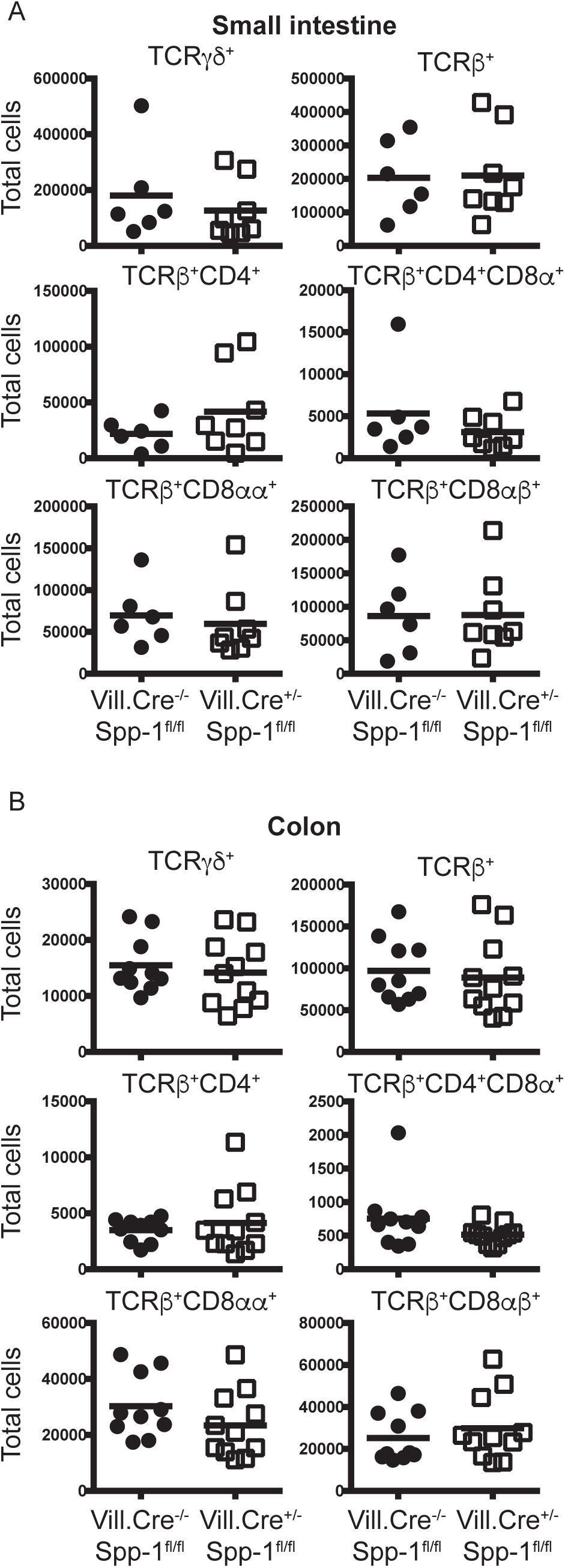
IEC-derived osteopontin does not promote IEL homeostasis. Small intestine (A) and colon (B) IEL were isolated from Villin-Cre^-/-^Spp-1^fl/fl^ and Villin-Cre^+/-^Spp-1^fl/fl^ mice, and the cellularity determined as indicated in Fig. 1A and B.

## Discussion

Intestinal IEL reside in the unique environment of the IEC monolayer. In this anatomical location, IEL are poised as one the first immunological defenses against potential pathogens from the intestinal lumen. In order to fulfill their immunological roles, IEL need to maintain their homeostasis. However, because IEL represent a diverse population of lymphoid cells, requirements for their homeostasis within the epithelium may depend on the particular type of IEL. For example, TCRγδ^+^ IEL require IL-7 for their proper development whereas other IEL are not affected by this cytokine (50). On the other hand, IL-15 deficiency does not disturb TCRγδ^+^ IEL but has a significant impact on TCRαβ^+^CD8αα^+^, iCD8α and iCD3^+^ IEL (7, 8, 51). The results presented in this report indicate that osteopontin is important for the homeostasis of many different IEL subpopulations. This implies that despite having different developmental pathways and cytokine requirements, the presence of osteopontin in the epithelium ensures proper homeostasis of most types of intestinal IEL.

Osteopontin-mediated T cell survival has been documented previously. For example, concanavalin A-activated T cells from lymph nodes show reduced levels of cell death in the presence of osteopontin (33). It is important to note that this report demonstrated a pivotal role for osteopontin as an enhancer for the survival of effector Th17 cells, particularly during brain inflammation (33). However, whereas this group studied differentiated Th17 cells in the context of brain inflammation our results are based on IEL in naïve animals. In the present report, using an *in vitro* system, we show that TCR^+^ IEL rapidly die in the absence of osteopontin, whereas the presence of this cytokine increased their *in vitro* survival (Fig. 3). Interestingly, each IEL subpopulation presented different survival kinetics, but appeared to have a similar survival requirement for osteopontin. Strikingly, the effect of osteopontin was only observed at 48 h (TCRβ^+^CD4^+^ and TCRβCD4^+^CD8α^+^ IEL) or 72 h (TCRγδ^+^and TCRβ^+^CD8α^+^ IEL). The reason why the osteopontin effect is only observed at later time points and not at 24 h is not clear. However, it is possible that as IEL are exposed to exogenous osteopontin during the first hours of culture, there is a decrease in their annexin V profile (as shown in Fig. 6), however their resilience to cell death is delayed, which is observed at later time points.

CD44 is known to be one of the receptors for osteopontin, and the interaction between these two molecules promotes adhesion and chemotaxis of bone marrow cells (52), increases growth of cancer cell lines (53), stimulates survival of pro-B cell lines (54), induces production of IL-10 in T cells (55), and protects T cell hybridomas from activated-induced cell death (56), among other functions. Despite the many roles attributed to the osteopontin-CD44 interaction, its role in IEL survival has not been investigated. IEL are considered to be in a “semi-activated” state (57), and most of these cells express CD44. Therefore, it is reasonable to speculate that some IEL subpopulations receive survival signals via the interaction between osteopontin and CD44. Interestingly, splenic CD44^+^ T cell survival was not increased by the addition of osteopontin *in vitro*, which suggests that osteopontin may not affect all T cells expressing CD44. We also show that the IEL survival promoted by osteopontin can be blunted by addition of anti-CD44 antibodies (Fig. 3), underscoring the importance of the osteopontin-CD44 interaction in IEL biology. However, it is noteworthy that IEL deficiencies in CD44^-/-^ mice do not faithfully resemble those observed in Spp-1^-/-^ mice, suggesting that in the absence of CD44, other receptors may bind osteopontin to stimulate survival of specific IEL subpopulations.

In the adoptive transfer experiments reported here, donor CD4 T cells from WT mice reconstituted the IEL compartment of Spp-1^-/-^Rag-2^-/-^ recipient mice less efficiently than in Rag-2^-/-^ recipient mice, suggesting that an environment capable of producing osteopontin is important for proper cell reconstitution in the intestines. However, transfer of T cells from osteopontin-deficient donor mice into Rag-2^-/-^ recipient mice resulted in reduced survival rates in the spleen and lymph nodes in comparison to donor T cells from wild type donor mice (31). These results indicate that intrinsic T cell-derived osteopontin is critical for normal cell reconstitution in secondary lymphoid organs whereas T cells present in the IEL compartment depend on osteopontin from the environment.

Adoptive transfer of total spleen T cells into immunodeficient hosts, such as Rag-2^-/-^ mice, normally results in cellular reconstitution and protection from T cell-mediated colitis due to the presence of regulatory T cells (58, 59). Surprisingly, when recipient Rag-2^-/-^ mice were deficient in osteopontin (Rag-2^-/-^Spp-1^-/-^ animals), mice developed colitis even in the presence of regulatory T cells (Fig. 4), raising the possibility that environmental osteopontin is important for maintaining regulatory T cell function, as shown in Fig. 5.

If most IEL subpopulations require osteopontin for their homeostasis, what are the cellular sources for this protein in the intestines? IEC produce osteopontin, and its expression increases during inflammation (25, 26). Other sources of osteopontin are within the IEL compartment, which appear to be confined to IEL expressing CD8α, including TCRγδ^+^, TCRβ^+^, and iCD8α cells (8, 35). We have previously shown that the survival of TCR^neg^ ILC1-like IEL (NKp46^+^NK1.1^+^) depends on iCD8α cell-derived osteopontin (36). Our previous publication and the results presented herein provide significance evidence to suggest that one of the main roles for iCD8α cells is the maintenance of proper IEL homeostasis. iCD8α cells constitute around 2 to 5% of the total IEL compartment. Considering that IEL are interspaced between IEC at an approximate ratio of 1 IEL for 10 IEC (2), how do iCD8α cells promote osteopontin-dependent survival of IEL in the monolayer of intestinal epithelial cells? It is known that some IEL, like TCRγδ^+^ cells scan the intestinal epithelium by moving out through the basal section of the epithelium and entering the IEC monolayer in another location (60). Therefore, it is possible that iCD8α cells use a similar mechanism to scan the monolayer, reach other IEL and provide needed osteopontin to other cells.

In the past few years, the role of osteopontin in the etiology of human diseases has been greatly appreciated. For example, recent work has investigated the use of neutralizing anti-osteopontin antibodies as a therapeutic with preclinical studies currently underway (ref. in (61)). Studies such as the one described herein show that osteopontin neutralization may affect IEL homeostasis in individuals with non-gastrointestinal tract disorders, but may be beneficial for IBD patients. Osteopontin appears to be a critical molecule with multiple effects, one of them supporting proper IEL homeostasis, and therefore additional studies are needed to better understand its function and how it affects the biology of the mucosal immune system.

## Acknowledgments

We thank the Flow Cytometry Shared Resource for technical help and guidance; the Translational Pathology Shared Resource for tissue processing; the Vanderbilt Genome Editing Resource, especially Dr. Leesa Sampson for her outstanding help for the generation of the Spp-1^fl/fl^ mice. We are thankful to Theresa Rogers, RN, for her help securing human blood and tissue samples. We thank Dr. Luc Van Kaer for reviewing the manuscript

## References

1. Cheroutre, H., F. Lambolez, and D. Mucida. 2011. The light and dark sides of intestinal intraepithelial lymphocytes. Nat Rev Immunol 11: 445–456.

2. Olivares-Villagomez, D., and L. Van Kaer. 2017. Intestinal Intraepithelial Lymphocytes: Sentinels of the Mucosal Barrier. Trends Immunol.

3. Van Kaer, L., and D. Olivares-Villagomez. 2018. Development, Homeostasis, and Functions of Intestinal Intraepithelial Lymphocytes. J Immunol 200: 2235–2244.

4. Fuchs, A., W. Vermi, J. S. Lee, S. Lonardi, S. Gilfillan, R. D. Newberry, M. Cella, and M. Colonna. 2013. Intraepithelial type 1 innate lymphoid cells are a unique subset of IL-12- and IL-15-responsive IFN-gamma-producing cells. Immunity 38: 769–781.

5. Talayero, P., E. Mancebo, J. Calvo-Pulido, S. Rodriguez-Munoz, I. Bernardo, R. Laguna-Goya, F. L. Cano-Romero, A. Garcia-Sesma, C. Loinaz, C. Jimenez, I. Justo, and E. Paz-Artal. 2016. Innate Lymphoid Cells Groups 1 and 3 in the Epithelial Compartment of Functional Human Intestinal Allografts. Am J Transplant 16: 72–82.

6. Van Acker, A., K. Gronke, A. Biswas, L. Martens, Y. Saeys, J. Filtjens, S. Taveirne, E. Van Ammel, T. Kerre, P. Matthys, T. Taghon, B. Vandekerckhove, J. Plum, I. R. Dunay, A. Diefenbach, and G. Leclercq. 2017. A Murine Intestinal Intraepithelial NKp46-Negative Innate Lymphoid Cell Population Characterized by Group 1 Properties. Cell Rep 19: 1431–1443.

7. Ettersperger, J., N. Montcuquet, G. Malamut, N. Guegan, S. Lopez-Lastra, S. Gayraud, C. Reimann, E. Vidal, N. Cagnard, P. Villarese, I. Andre-Schmutz, R. Gomes Domingues, C. Godinho-Silva, H. Veiga-Fernandes, L. Lhermitte, V. Asnafi, E. Macintyre, C. Cellier, K. Beldjord, J. P. Di Santo, N. Cerf-Bensussan, and B. Meresse. 2016. Interleukin-15-Dependent T-Cell-like Innate Intraepithelial Lymphocytes Develop in the Intestine and Transform into Lymphomas in Celiac Disease. Immunity 45: 610–625.

8. Van Kaer, L., H. M. Algood, K. Singh, V. V. Parekh, M. J. Greer, M. B. Piazuelo, J. H. Weitkamp, P. Matta, R. Chaturvedi, K. T. Wilson, and D. Olivares-Villagomez. 2014. CD8alphaalpha(+) Innate-Type Lymphocytes in the Intestinal Epithelium Mediate Mucosal Immunity. Immunity 41: 451–464.

9. Edelblum, K. L., G. Singh, M. A. Odenwald, A. Lingaraju, K. El Bissati, R. McLeod, A. I. Sperling, and J. R. Turner. 2015. gammadelta Intraepithelial Lymphocyte Migration Limits Transepithelial Pathogen Invasion and Systemic Disease in Mice. Gastroenterology 148: 1417–1426.

10. Ismail, A. S., K. M. Severson, S. Vaishnava, C. L. Behrendt, X. Yu, J. L. Benjamin, K. A. Ruhn, B. Hou, A. L. DeFranco, F. Yarovinsky, and L. V. Hooper. 2011. Gammadelta intraepithelial lymphocytes are essential mediators of host-microbial homeostasis at the intestinal mucosal surface. Proc Natl Acad Sci U S A 108: 8743–8748.

11. Inagaki-Ohara, K., T. Chinen, G. Matsuzaki, A. Sasaki, Y. Sakamoto, K. Hiromatsu, F. Nakamura-Uchiyama, Y. Nawa, and A. Yoshimura. 2004. Mucosal T cells bearing TCRgammadelta play a protective role in intestinal inflammation. J Immunol 173: 1390–1398.

12. Lepage, A. C., D. Buzoni-Gatel, D. T. Bout, and L. H. Kasper. 1998. Gut-derived intraepithelial lymphocytes induce long term immunity against Toxoplasma gondii. J Immunol 161: 4902–4908.

13. Masopust, D., D. Choo, V. Vezys, E. J. Wherry, J. Duraiswamy, R. Akondy, J. Wang, K. A. Casey, D. L. Barber, K. S. Kawamura, K. A. Fraser, R. J. Webby, V. Brinkmann, E. C. Butcher, K. A. Newell, and R. Ahmed. 2010. Dynamic T cell migration program provides resident memory within intestinal epithelium. J Exp Med 207: 553–564.

14. Masopust, D., J. Jiang, H. Shen, and L. Lefrancois. 2001. Direct analysis of the dynamics of the intestinal mucosa CD8 T cell response to systemic virus infection. J Immunol 166: 2348–2356.

15. Das, G., M. M. Augustine, J. Das, K. Bottomly, P. Ray, and A. Ray. 2003. An important regulatory role for CD4+CD8 alpha alpha T cells in the intestinal epithelial layer in the prevention of inflammatory bowel disease. Proc Natl Acad Sci U S A 100: 5324-5329. PMC1535703.

16. Kumar, A. A., A. G. Delgado, M. B. Piazuelo, L. Van Kaer, and D. Olivares-Villagomez. 2017. Innate CD8alphaalpha(+) lymphocytes enhance anti-CD40 antibody-mediated colitis in mice. Immun Inflamm Dis 5: 109–123.

17. Franzen, A., and D. Heinegard. 1985. Isolation and characterization of two sialoproteins present only in bone calcified matrix. Biochem J 232: 715–724.

18. Prince, C. W., T. Oosawa, W. T. Butler, M. Tomana, A. S. Bhown, M. Bhown, and R. E. Schrohenloher. 1987. Isolation, characterization, and biosynthesis of a phosphorylated glycoprotein from rat bone. J Biol Chem 262: 2900–2907.

19. Di Bartolomeo, M., F. Pietrantonio, A. Pellegrinelli, A. Martinetti, L. Mariani, M. G. Daidone, E. Bajetta, G. Pelosi, F. de Braud, I. Floriani, and R. Miceli. 2016. Osteopontin, E-cadherin, and beta-catenin expression as prognostic biomarkers in patients with radically resected gastric cancer. Gastric Cancer 19: 412–420.

20. Giachelli, C. M., N. Bae, M. Almeida, D. T. Denhardt, C. E. Alpers, and S. M. Schwartz. 1993. Osteopontin is elevated during neointima formation in rat arteries and is a novel component of human atherosclerotic plaques. J Clin Invest 92: 1686–1696.

21. Icer, M. A., and M. Gezmen-Karadag. 2018. The multiple functions and mechanisms of osteopontin. Clin Biochem 59: 17–24.

22. Oz, H. S., J. Zhong, and W. J. de Villiers. 2012. Osteopontin ablation attenuates progression of colitis in TNBS model. Dig Dis Sci 57: 1554–1561.

23. Zhong, J., E. R. Eckhardt, H. S. Oz, D. Bruemmer, and W. J. de Villiers. 2006. Osteopontin deficiency protects mice from Dextran sodium sulfate-induced colitis. Inflamm Bowel Dis 12: 790–796.

24. Mishima, R., F. Takeshima, T. Sawai, K. Ohba, K. Ohnita, H. Isomoto, K. Omagari, Y. Mizuta, Y. Ozono, and S. Kohno. 2007. High plasma osteopontin levels in patients with inflammatory bowel disease. J Clin Gastroenterol 41: 167–172.

25. Sato, T., T. Nakai, N. Tamura, S. Okamoto, K. Matsuoka, A. Sakuraba, T. Fukushima, T. Uede, and T. Hibi. 2005. Osteopontin/Eta-1 upregulated in Crohn’s disease regulates the Th1 immune response. Gut 54: 1254–1262.

26. Gassler, N., F. Autschbach, S. Gauer, J. Bohn, B. Sido, H. F. Otto, H. Geiger, and N. Obermuller. 2002. Expression of osteopontin (Eta-1) in Crohn disease of the terminal ileum. Scand J Gastroenterol 37: 1286–1295.

27. Masuda, H., Y. Takahashi, S. Asai, and T. Takayama. 2003. Distinct gene expression of osteopontin in patients with ulcerative colitis. J Surg Res 111: 85–90.

28. Neuman, M. G. 2012. Osteopontin biomarker in inflammatory bowel disease, animal models and target for drug discovery. Dig Dis Sci 57: 1430–1431.

29. Boumans, M. J., J. G. Houbiers, P. Verschueren, H. Ishikura, R. Westhovens, E. Brouwer, B. Rojkovich, S. Kelly, M. den Adel, J. Isaacs, H. Jacobs, J. Gomez-Reino, G. M. Holtkamp, A. Hastings, D. M. Gerlag, and P. P. Tak. 2012. Safety, tolerability, pharmacokinetics, pharmacodynamics and efficacy of the monoclonal antibody ASK8007 blocking osteopontin in patients with rheumatoid arthritis: a randomised, placebo controlled, proof-of-concept study. Annals of the rheumatic diseases 71: 180–185.

30. Lund, S. A., C. L. Wilson, E. W. Raines, J. Tang, C. M. Giachelli, and M. Scatena. 2013. Osteopontin mediates macrophage chemotaxis via alpha4 and alpha9 integrins and survival via the alpha4 integrin. J Cell Biochem 114: 1194–1202.

31. Leavenworth, J. W., B. Verbinnen, Q. Wang, E. Shen, and H. Cantor. 2015. Intracellular osteopontin regulates homeostasis and function of natural killer cells. Proc Natl Acad Sci U S A 112: 494–499.

32. Shinohara, M. L., H. J. Kim, J. H. Kim, V. A. Garcia, and H. Cantor. 2008. Alternative translation of osteopontin generates intracellular and secreted isoforms that mediate distinct biological activities in dendritic cells. Proc Natl Acad Sci U S A 105: 7235–7239.

33. Hur, E. M., S. Youssef, M. E. Haws, S. Y. Zhang, R. A. Sobel, and L. Steinman. 2007. Osteopontin-induced relapse and progression of autoimmune brain disease through enhanced survival of activated T cells. Nat Immunol 8: 74–83.

34. Ashkar, S., G. F. Weber, V. Panoutsakopoulou, M. E. Sanchirico, M. Jansson, S. Zawaideh, S. R. Rittling, D. T. Denhardt, M. J. Glimcher, and H. Cantor. 2000. Eta-1 (osteopontin): an early component of type-1 (cell-mediated) immunity. Science 287: 860–864.

35. Ito, K., A. Nakajima, Y. Fukushima, K. Suzuki, K. Sakamoto, Y. Hamazaki, K. Ogasawara, N. Minato, and M. Hattori. 2017. The potential role of Osteopontin in the maintenance of commensal bacteria homeostasis in the intestine. PLoS One 12: e0173629.

36. Nazmi, A., K. L. Hoek, M. J. Greer, M. B. Piazuelo, N. Minato, and D. Olivares-Villagomez. 2019. Innate CD8alphaalpha+ cells promote ILC1-like intraepithelial lymphocyte homeostasis and intestinal inflammation. PLoS One 14: e0215883.

37. Quadros, R. M., H. Miura, D. W. Harms, H. Akatsuka, T. Sato, T. Aida, R. Redder, G. P. Richardson, Y. Inagaki, D. Sakai, S. M. Buckley, P. Seshacharyulu, S. K. Batra, M. A. Behlke, S. A. Zeiner, A. M. Jacobi, Y. Izu, W. B. Thoreson, L. D. Urness, S. L. Mansour, M. Ohtsuka, and C. B. Gurumurthy. 2017. Easi-CRISPR: a robust method for one-step generation of mice carrying conditional and insertion alleles using long ssDNA donors and CRISPR ribonucleoproteins. Genome biology 18: 92.

38. Olivares-Villagomez, D., Y. V. Mendez-Fernandez, V. V. Parekh, S. Lalani, T. L. Vincent, H. Cheroutre, and L. Van Kaer. 2008. Thymus leukemia antigen controls intraepithelial lymphocyte function and inflammatory bowel disease. Proc Natl Acad Sci U S A 105: 17931-17936. PMC2584730.

39. Hoek, K. L., P. Samir, L. M. Howard, X. Niu, N. Prasad, A. Galassie, Q. Liu, T. M. Allos, K. A. Floyd, Y. Guo, Y. Shyr, S. E. Levy, S. Joyce, K. M. Edwards, and A. J. Link. 2015. A cell-based systems biology assessment of human blood to monitor immune responses after influenza vaccination. PLoS One 10: e0118528.

40. Weitkamp, J. H., T. Koyama, M. T. Rock, H. Correa, J. A. Goettel, P. Matta, K. Oswald-Richter, M. J. Rosen, B. G. Engelhardt, D. J. Moore, and D. B. Polk. 2013. Necrotising enterocolitis is characterised by disrupted immune regulation and diminished mucosal regulatory (FOXP3)/effector (CD4, CD8) T cell ratios. Gut 62: 73–82.

41. Aranda, R., B. C. Sydora, P. L. McAllister, S. W. Binder, H. Y. Yang, S. R. Targan, and M. Kronenberg. 1997. Analysis of intestinal lymphocytes in mouse colitis mediated by transfer of CD4+, CD45RBhigh T cells to SCID recipients. J Immunol 158: 3464–3473.

42. Patro, R., G. Duggal, M. I. Love, R. A. Irizarry, and C. Kingsford. 2017. Salmon provides fast and bias-aware quantification of transcript expression. Nature methods 14: 417–419.

43. Robinson, M. D., D. J. McCarthy, and G. K. Smyth. 2010. edgeR: a Bioconductor package for differential expression analysis of digital gene expression data. Bioinformatics 26: 139–140.

44. Subramanian, A., P. Tamayo, V. K. Mootha, S. Mukherjee, B. L. Ebert, M. A. Gillette, A. Paulovich, S. L. Pomeroy, T. R. Golub, E. S. Lander, and J. P. Mesirov. 2005. Gene set enrichment analysis: a knowledge-based approach for interpreting genome-wide expression profiles. Proc Natl Acad Sci U S A 102: 15545–15550.

45. Yokosaki, Y., K. Tanaka, F. Higashikawa, K. Yamashita, and A. Eboshida. 2005. Distinct structural requirements for binding of the integrins alphavbeta6, alphavbeta3, alphavbeta5, alpha5beta1 and alpha9beta1 to osteopontin. Matrix biology : journal of the International Society for Matrix Biology 24: 418–427.

46. Katagiri, Y. U., J. Sleeman, H. Fujii, P. Herrlich, H. Hotta, K. Tanaka, S. Chikuma, H. Yagita, K. Okumura, M. Murakami, I. Saiki, A. F. Chambers, and T. Uede. 1999. CD44 variants but not CD44s cooperate with beta1-containing integrins to permit cells to bind to osteopontin independently of arginine-glycine-aspartic acid, thereby stimulating cell motility and chemotaxis. Cancer Res 59: 219–226.

47. Beura, L. K., S. E. Hamilton, K. Bi, J. M. Schenkel, O. A. Odumade, K. A. Casey, E. A. Thompson, K. A. Fraser, P. C. Rosato, A. Filali-Mouhim, R. P. Sekaly, M. K. Jenkins, V. Vezys, W. N. Haining, S. C. Jameson, and D. Masopust. 2016. Normalizing the environment recapitulates adult human immune traits in laboratory mice. Nature 532: 512–516.

48. Smolewski, P., and T. Robak. 2011. Inhibitors of apoptosis proteins (IAPs) as potential molecular targets for therapy of hematological malignancies. Curr Mol Med 11: 633–649.

49. Bollyky, P. L., B. A. Falk, S. A. Long, A. Preisinger, K. R. Braun, R. P. Wu, S. P. Evanko, J. H. Buckner, T. N. Wight, and G. T. Nepom. 2009. CD44 costimulation promotes FoxP3+ regulatory T cell persistence and function via production of IL-2, IL-10, and TGF-beta. J Immunol 183: 2232–2241.

50. Maki, K., S. Sunaga, Y. Komagata, Y. Kodaira, A. Mabuchi, H. Karasuyama, K. Yokomuro, J. I. Miyazaki, and K. Ikuta. 1996. Interleukin 7 receptor-deficient mice lack gammadelta T cells. Proc Natl Acad Sci U S A 93: 7172–7177. PMC38955.

51. Kennedy, M. K., M. Glaccum, S. N. Brown, E. A. Butz, J. L. Viney, M. Embers, N. Matsuki, K. Charrier, L. Sedger, C. R. Willis, K. Brasel, P. J. Morrissey, K. Stocking, J. C. Schuh, S. Joyce, and J. J. Peschon. 2000. Reversible defects in natural killer and memory CD8 T cell lineages in interleukin 15-deficient mice. J Exp Med 191: 771–780.

52. Weber, G. F., S. Ashkar, M. J. Glimcher, and H. Cantor. 1996. Receptor-ligand interaction between CD44 and osteopontin (Eta-1). Science 271: 509–512.

53. Sun, S. J., C. C. Wu, G. T. Sheu, H. Y. Chang, M. Y. Chen, Y. Y. Lin, C. Y. Chuang, S. L. Hsu, and J. T. Chang. 2016. Integrin beta3 and CD44 levels determine the effects of the OPN-a splicing variant on lung cancer cell growth. Oncotarget 7: 55572–55584.

54. Lin, Y. H., and H. F. Yang-Yen. 2001. The osteopontin-CD44 survival signal involves activation of the phosphatidylinositol 3-kinase/Akt signaling pathway. J Biol Chem 276: 46024–46030.

55. Murugaiyan, G., A. Mittal, and H. L. Weiner. 2008. Increased osteopontin expression in dendritic cells amplifies IL-17 production by CD4+ T cells in experimental autoimmune encephalomyelitis and in multiple sclerosis. J Immunol 181: 7480–7488.

56. Larkin, J., G. J. Renukaradhya, V. Sriram, W. Du, J. Gervay-Hague, and R. R. Brutkiewicz. 2006. CD44 differentially activates mouse NK T cells and conventional T cells. J Immunol 177: 268–279.

57. Montufar-Solis, D., T. Garza, and J. R. Klein. 2007. T-cell activation in the intestinal mucosa. Immunol Rev 215: 189-201. PMC2754816.

58. Powrie, F., R. Correa-Oliveira, S. Mauze, and R. L. Coffman. 1994. Regulatory interactions between CD45RBhigh and CD45RBlow CD4+ T cells are important for the balance between protective and pathogenic cell-mediated immunity. J Exp Med 179: 589-600. PMC2191378.

59. Powrie, F., M. W. Leach, S. Mauze, S. Menon, L. B. Caddle, and R. L. Coffman. 1994. Inhibition of Th1 responses prevents inflammatory bowel disease in scid mice reconstituted with CD45RBhi CD4+ T cells. Immunity 1: 553–562.

60. Hoytema van Konijnenburg, D. P., B. S. Reis, V. A. Pedicord, J. Farache, G. D. Victora, and D. Mucida. 2017. Intestinal Epithelial and Intraepithelial T Cell Crosstalk Mediates a Dynamic Response to Infection. Cell 171: 783–794 e713.

61. Farrokhi, V., J. R. Chabot, H. Neubert, and Z. Yang. 2018. Assessing the Feasibility of Neutralizing Osteopontin with Various Therapeutic Antibody Modalities. Sci Rep 8: 7781.

