## Supplemental Figures 1 to 4 for "Osteopontin and iCD8α cells promote intestinal intraepithelial lymphocyte homeostasis"

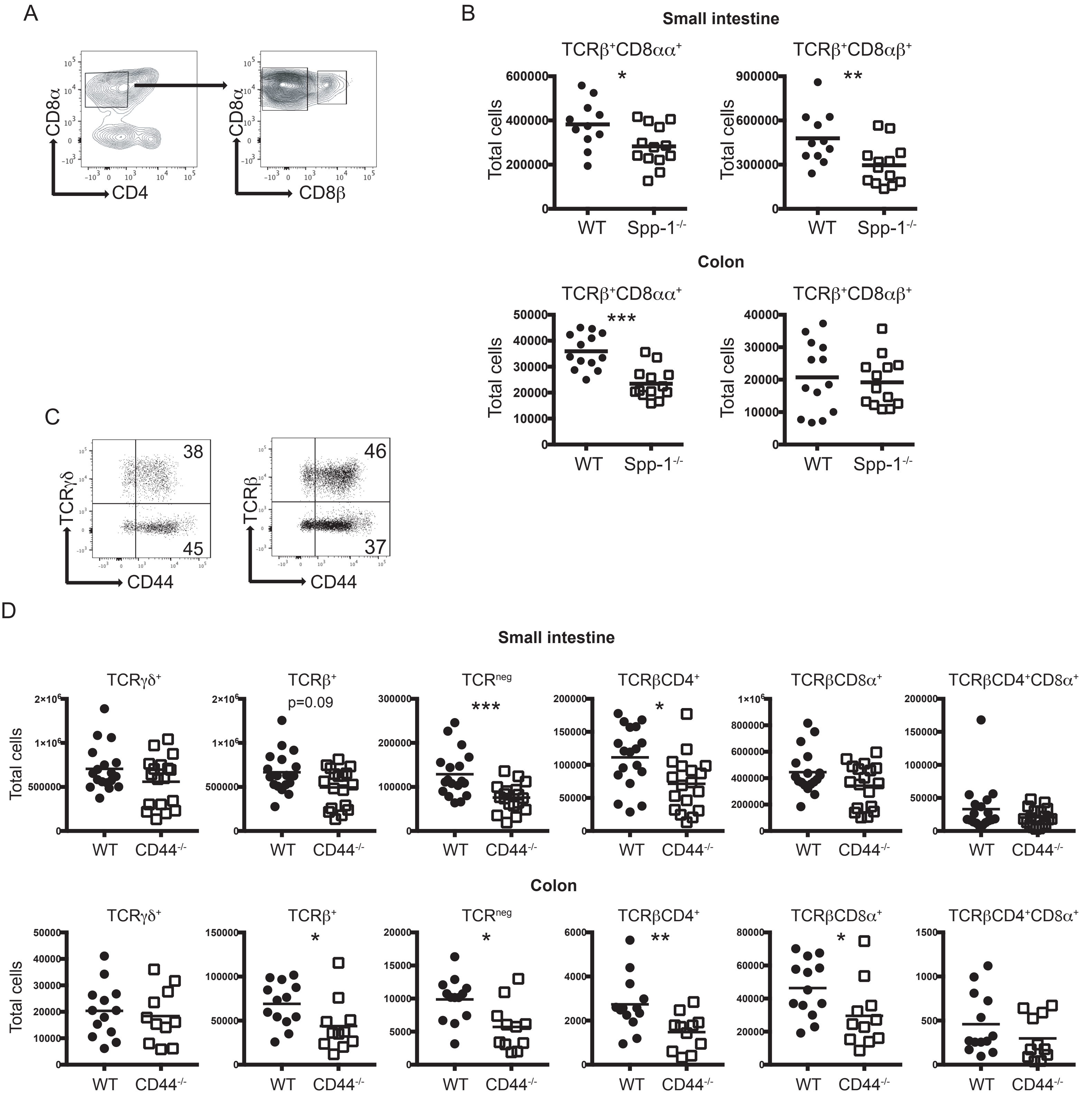


Supplemental Figure 1. (A) Small intestine and colon IEL were analyzed as in Figure 1A. CD8α$^{+}$ IEL were subsequently divided by CD8β expression. (B) Total IEL numbers from the small intestine (top) and colons (bottom) of WT and Spp-1$^{-/-}$ mice. Data is combined of three independent experiments. Each dot represents an individual sample (n = 11 to 13). (C) CD44 expression in TCRγδ$^{+}$ and TCRβ$^{+}$IEL from WT mice. After excluding dead cells and IEC, cells were gated as CD45$^{+}$ cells. (D) Total number of different IEL subpopulations derived from the small intestine of colon from WT and CD44$^{-/-}$ mice. Data is combined of three independent experiments. Each dot represents an individual sample (n = 12 to 19).*P<0.05,**P<0.01,***P<0.001.

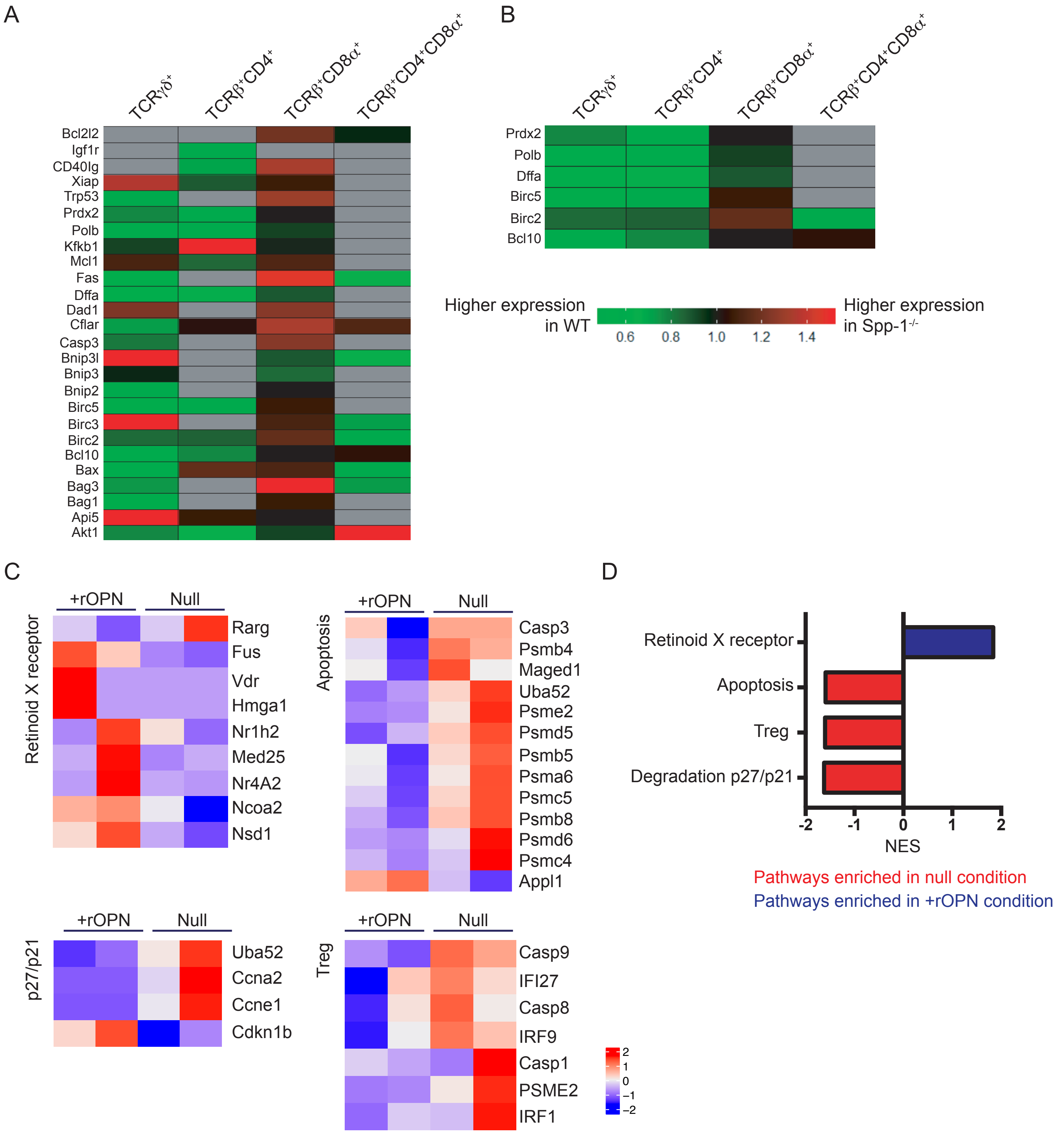


Supplemental Figure 2. Osteopontin induces a survival program in IEL. (A) RNA was isolated from FACS-enriched TCRγδ$^+$, TCRβ$^+$CD4$^+$, TCRβ$^+$CD8α$^+$ and TCRβ$^+$CD4$^+$CD8α$^+$ IEL from WT and Spp-1$^{-/-}$ mice, and used to detect expression of anti-apoptosis-related genes using a QIAGEN RT2 Profiler PCR Array. Each column represents the average of 4 samples from individual mice. Heat-map is a representative experiment of two performed. (B) Heat-map representing a sample of genes differentially expressed in most WT IEL populations; data were derived from (A). (C) Enriched CD45$^+$ IEL derived from 2 individual WT mice were pooled into a biological replicate. A total of 8 mice were used yielding 4 replicates. Half the cells from each replicate were incubated in the presence or absence of osteopontin (2 mg/ml) for 24h. RNA-seq was used to detect gene expression profile. Each column depicts a representative biological replicate. (D) Gene set enrichment analysis summary.

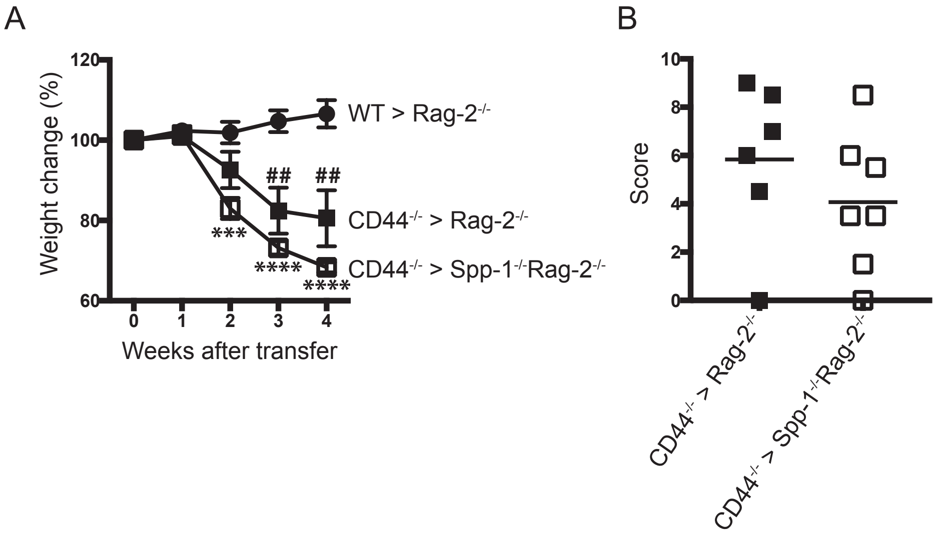


Supplemental Figure 3. (A) Enriched total T cells from CD44$^{-/-}$ mice were adoptively transferred into Rag-2$^{-/-}$ and Spp-1$^{-/-}$Rag-2$^{-/-}$ mice, and weights were monitored weekly. (B) Colon pathology scored as described in Fig. 4D. Data are representative of two independent experiments. Each dot represents an individual sample (n=6 to 7). ***$P$<0.001 and ****$P$<0.0001 between WT > Rag-2$^{-/-}$ and CD44 > Spp-1$^{-/-}$Rag-2$^{-/-}$ groups; $^{\#\#}P$<0.01 between WT > Rag-2$^{-/-}$ and CD44 > Rag-2$^{-/-}$ groups (Mann-Whitney U test).

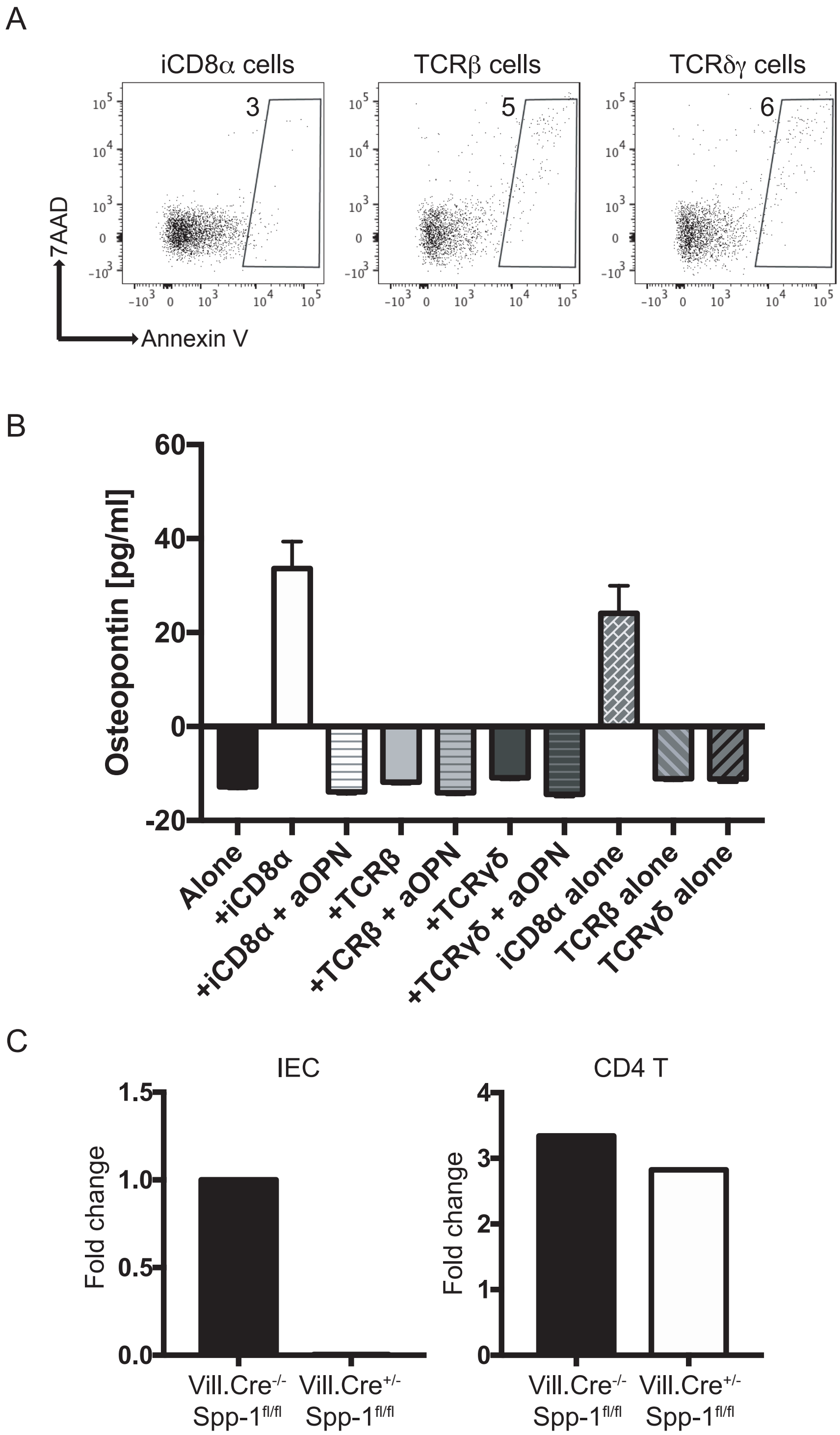


Supplemental Figure 4. (A) Viability of osteopontin-competent cells after 4 h of culture. Cells were analyzed as indicated in Fig. 6. (B) $CD45^+$ IEL from Spp-$1^{-/-}$ mice were cultured in the presence or absence of the indicated osteopontin-competent IEL populations, with or without anti-osteopontin antibodies. After 4 h of culture, supernatants were recovered and osteopontin concentration measured by ELISA. Results are from two pooled independent experiments. The last three bars indicate culture of the osteopontin-competent IEL poppulations without IEL from Spp-$1^{-/-}$ mice. (C) Real-time PCR analysis from total RNA isolated from enriched colon IEC ($CD45^{neg}$ $g8.8^+$ cells) (left) from a pool of 3 mice treated with 2.5% DSS in the drinking water for 5 days, cells were sorted 2 days after DSS treatment; or anti-CD3/CD28 activated spleen CD4 T cells (right) from the indicated mice.
